# Individualized ctDNA Fingerprints to Monitor Treatment Response and Recurrence in Multiple Cancer Types

**DOI:** 10.1101/732503

**Authors:** Jiaping Li, Wei Jiang, Jinwang Wei, Jianwei Zhang, Linbo Cai, Minjie Luo, Zhan Wang, Wending Sun, Chen Wang, Chun Dai, Guan Wang, Qiang Xu, Yanhong Deng

## Abstract

Circulating tumor DNA (ctDNA) panels hold high promise of accurately predicting the therapeutic response of tumors while being minimally invasive and cost-efficient. However, their use has been limited to a small number of tumor types and patients. Here, we developed individualized ctDNA fingerprints suitable for most patients with multiple cancer types. The panels were designed based on individual whole-exome sequencing data in 521 Chinese patients and targeting high clonal population clusters of somatic mutations. Together, these patients represent 12 types of cancers and seven different treatments. The customized ctDNA panels have a median somatic mutation number of 19, most of which are patient-specific rather than cancer hotspot mutations; 66.8% of the patients were ctDNA-positive. We further evaluated the ctDNA content fraction (CCF) of the mutations, and analyzed the association between the change of ctDNA concentration and therapeutic response. We followed up 106 patients for clinical evaluation, demonstrating a significant correlation of changes in ctDNA with clinical outcomes, with a consistency rate of 93.4%. In particular, the median CCF increased by 204.6% in patients with progressive disease, decreased by 82.5% in patients with remission, and was relatively stable in patients with stable disease. Overall, 85% of the patients with a ctDNA-positive status experienced metastasis or relapse long before imaging detection, except for two patients who developed recurrence and metastasis almost simultaneously. The average lead time between the first ctDNA-positive finding and radiological diagnosis was 76 days in three patients that changed from a ctDNA-negative to -positive status. Our individualized ctDNA analysis can effectively monitor the treatment response, metastasis, and recurrence in multiple cancer types in patients with multiple treatment options, therefore offering great clinical applicability for improving personalized treatment in cancer.

**One Sentence Summary:** ctDNA fingerprint panels were customized to predict the treatment response for multiple cancer types from individual whole-exome sequencing data.

## Introduction

The incidence of cancer is increasing worldwide (1). Despite continuous development of new treatment strategies, the therapeutic effectiveness and recurrence remain difficult to monitor and it is hard to predict disease progress, which is essential for making appropriate clinical decisions. Although radiological imaging and protein biomarker analysis in the blood remain the most common methods of predicting treatment outcome in cancer, they both lack sensitivity and specificity (2, 3). Thus, new methods with high sensitivity and specificity are needed to monitor the treatment response in real-time and assist in the timely adjustment of a therapeutic regimen according to the most recent gene mutation status.

Circulating tumor DNA (ctDNA) has emerged as a new biomarker to assess therapeutic efficacy and detect recurrence. ctDNA is derived from apoptotic or necrotic tumor cells and resides in the blood, allowing for repeatable sampling in a simple and non-invasive manner. Importantly, analysis of ctDNA can provide snapshots of the tumor burden and genomic profile because ctDNA has a short half-life and is highly specific to the tumor from which it is derived (4). Indeed, several studies have demonstrated the value of ctDNA in monitoring the treatment response. In a study of 53 patients with metastatic colorectal cancer (CRC) who received standard first-line chemotherapy, the fold change of the ctDNA level was correlated with the tumor response exhibited by computed tomography (CT) imaging at 8–10 weeks post-treatment (5). In another study, the majority of 46 patients who were initially diagnosed as having non-metastatic triple-negative breast cancer showed metastasis and recurrence after one cycle of neoadjuvant chemotherapy, and all of these metastatic patients were found to be ctDNA-positive (6). In an open phase I clinical trial of the combination of BRAF, EGFR, and MEK inhibitors, patients with the BRAF V600E mutation showed a greater change of ctDNA and a higher ratio of partial response (PR) and complete response (CR) to treatments than the patients with less of a change of ctDNA and showing stable disease (SD) or progressive disease (PD) status. Notably, there was no significant difference in the expression level of the common protein biomarker carcinoembryonic antigen (CEA) between these groups (7). Therefore, ctDNA appears to be superior to standard radiological and protein tumor markers in predicting the therapeutic response.

In contrast to generic panels containing only tumor hotspot mutations, a ctDNA panel can be tailored to the tumor genetic profile of individual patients. The TRAcking non-small cell lung cancer evolution through therapy (Rx) (TRACERx) study customized each patient’s ctDNA panel based on clonal mutations, which included a median of 11 clonal and six subclonal single-nucleotide variants (SNVs) per patient (8). The TRACERx study demonstrated that the tumor volume correlated with the allele variation frequency of the average clonal SNVs, and that dynamic measurement of plasma ctDNA could predict and confirm tumor recurrence earlier than clinical CT imaging (8). In another study, a ctDNA panel with 9–24 founding mutations detected residual disease in five out of the six patients with breast cancer after adjuvant therapy; the failure in detecting residual disease for the last patient was attributed to an insufficient blood sample for ctDNA analysis (9). These studies highlighted the successful use of customized ctDNA panels in monitoring lung and breast cancers.

Here, we developed a new platform of individualized ctDNA panels designed based on whole-exome sequencing (WES) data of individual patients. We used SciClone tools (10) to calculate and select the top 20–40 somatic mutations within high clonal population clusters to custom individualized ctDNA panels, and the ctDNA content fraction (CCF) was estimated based on the proportion of these acellular tumor somatic mutant DNA fragments in total cell-free DNA. We also used a pre-designed ctDNA panel with eight hotspot cancer genes to monitor acquired resistance mutations for comparison. We applied the new platform to detect ctDNA in longitudinally collected plasma samples from 521 Chinese patients who represented more than 12 tumor types and had received seven types of treatments, and explored the ability of using ctDNA to predict the treatment response and recurrence.

## Results

### Validation of the individualized ctDNA fingerprint assay in reference and tumor samples

To determine the detection threshold and sensitivity of the individualized ctDNA assay, we performed more than 150 tests for each variant with a gradient of allelic frequencies in the range of 0–15%. Ten nanograms of ctDNA was used for each test. The background ctDNA value in the wild-type reference samples was 0.065% ± 0.062% (mean ± standard deviation). We set the sum of the mean and standard deviation, 0.127%, as the threshold, which represents the upper confidence level of background noise. The specificity of detecting an individual mutation in the wild-type samples was 80.3%, and the sensitivities of reference samples with expected allele frequencies of 0.1%, 0.25%, and 0.5% were 40.6%, 75.0%, and 96.3%, respectively. Mutations with a known frequency greater than 1% were all successfully detected (Supplementary Table S1). Since each allele has two copies, the threshold CCF for positive detection is twice the reference sample threshold at about 0.25%.

In addition to the sensitivity of a single variant, we calculated the sensitivities of a panel containing five and 10 mutants at concentrations of 0.10%, 0.25%, and 0.50% standard references, which were 44.4% and 46.0%, 97.6% and 99.8%, and 99.99% and 100%, respectively. The specificities of the five and 10 mutants in wild-type samples were 98.7% and 99.9% respectively. Therefore, a panel with more tumor-specific SNVs shows higher sensitivity and specificity. Based on these results, patients with CCF values higher than 0.25% were considered to be ctDNA-positive, although CCF values were still recorded for those lower than 0.25%.

### Patients and study design

Among the 552 cancer patients recruited in this study, WES data were obtained from 521 patients who had case-matched tumor specimen and blood samples. Individualized ctDNA assays were designed for each patient based on WES data using a median of 26 pairs of primers, ranging from 7 to 39. After filtering out amplicons with low amplification efficiency and poor quality, the number of amplicons included in the individualized ctDNA fingerprint assays ranged from 4 to 33, with a median of 19. All patients underwent at least two independent ctDNA tests, with some patients undergoing more than three tests. In total, 1436 individualized ctDNA tests were performed on the 521 patients. Among them, 106 patients received follow-up clinical evaluations: 80 patients had one follow-up visit, 21 patients had two follow-up visits, and five patients had three or more follow-up visits. The flow of the study design is illustrated in Figure 1.

**Fig. 1.**
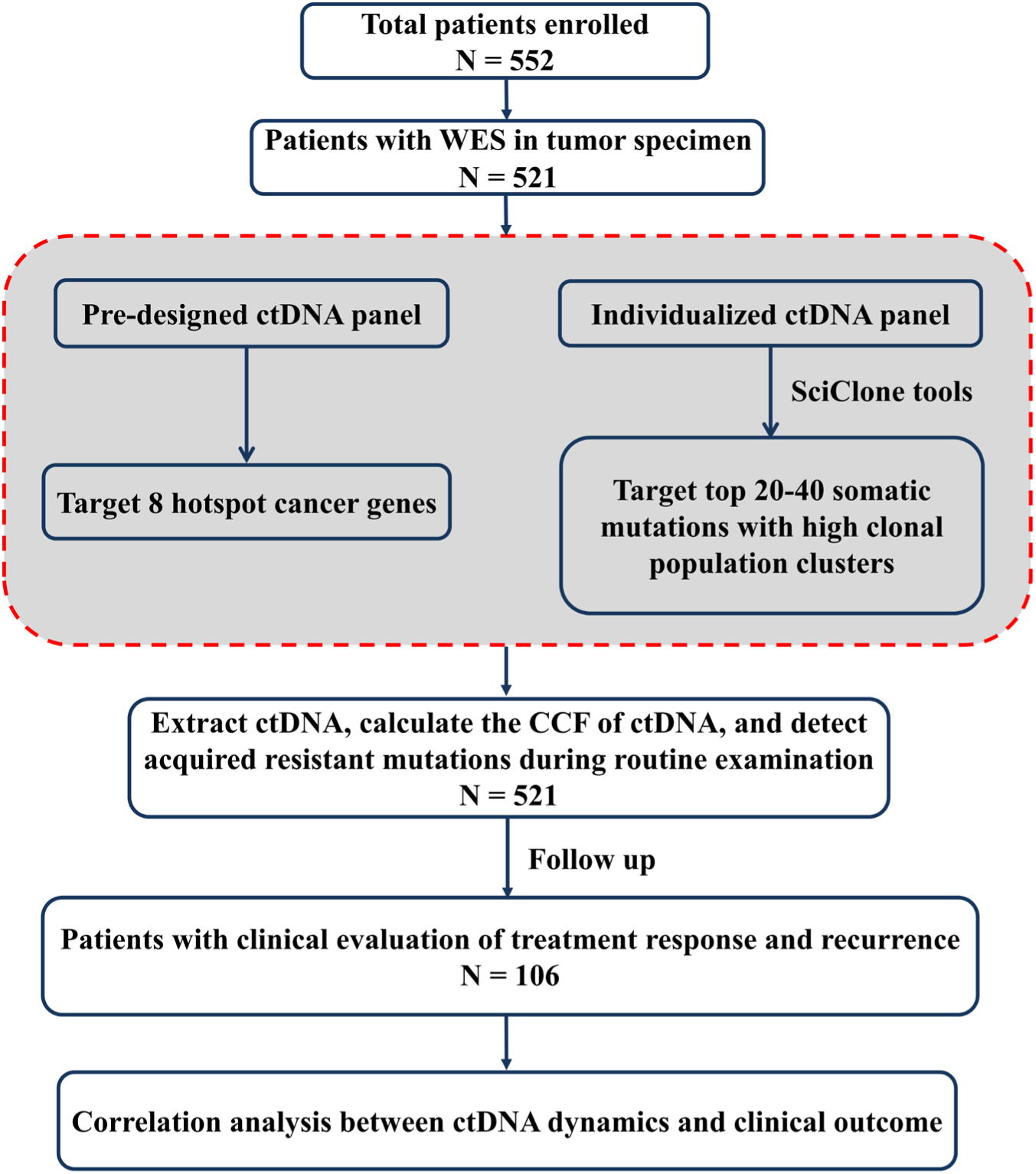
Patient characterization and flow chart of the study design. The dotted red box shows the pre-designed ctDNA panel and individualized ctDNA panel design.

### Application of individualized ctDNA fingerprint assays across multiple tumor types

The distribution of tumor types across the 521 Chinese patients is shown in Figure 2A. The highest number of patients had colorectal cancer (CRC), followed by liver cancer, glioma [combined result of blood and cerebrospinal fluid (CSF) tests], and lung cancer in significant numbers; other cancer types included gastric cancer, cholangiocarcinoma, and pancreatic cancer. For the 15 patients with glioma, we additionally collected CSF along with the total 70 plasma samples. Overall, 66.8% of the patients were positive for ctDNA, and 50.9%, 43.0%, and 25.7% of the patients had detectable ctDNA with a CCF above 0.5%, 1%, and 5% respectively. In the glioma samples, ctDNA detection in the blood was significantly lower than that in the CSF (24.3% vs. 80.0%, p < 0.01; Figure 2B), presumably due to blockage of the blood-brain barrier (BBB).

**Fig. 2.**
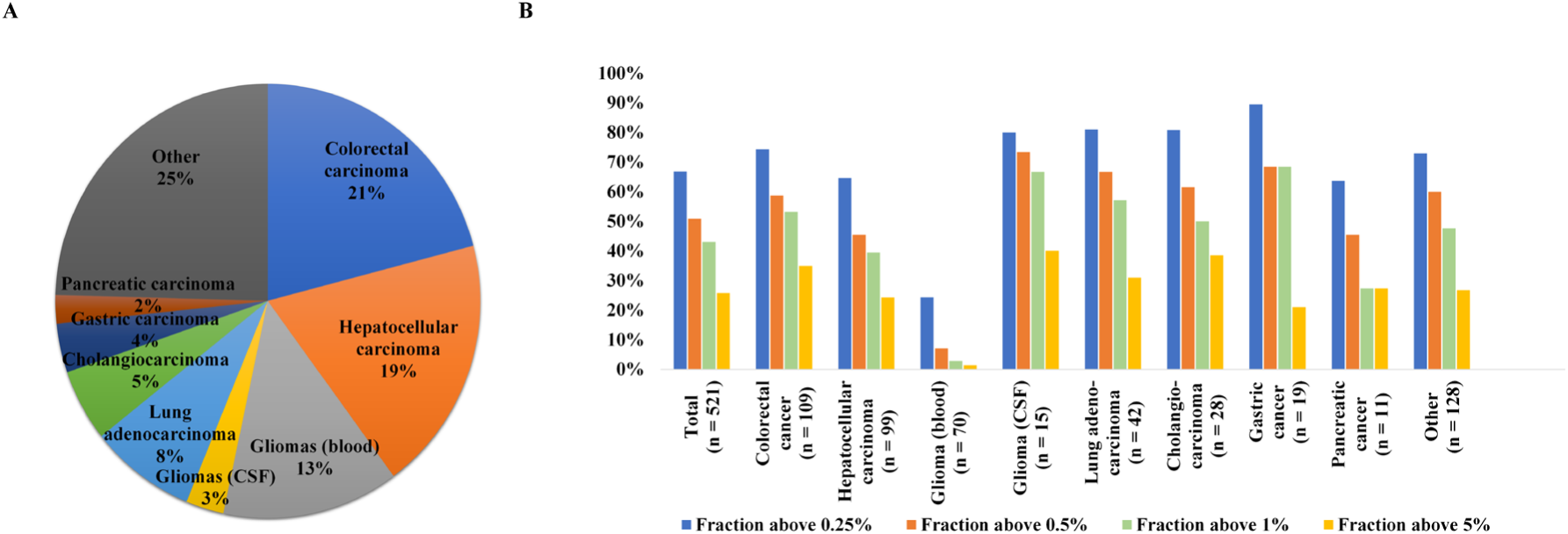
Overview of ctDNA detection. (A) Distribution of tumor types in the 521 patients. (B) Fraction of patients with a ctDNA detection rate above 0.25% (blue), 0.5% (orange), 1% (light green), and 5% (yellow) in different cancer types.

We also tested for acquired drug-related mutations in 16 patients using a pre-designed ctDNA panel with eight hotspot cancer genes (BRAF, EGFR, ERBB2, KIT, KRAS, MET, NRAS, and PIK3CA). The majority of these patients had colon (n = 6) and lung (n = 3) cancers: all CRC patients acquired a KRAS mutation with a frequency >0.05%; two of the lung adenocarcinoma patients acquired the *EGFR* p.T790M mutation, one with a frequency of 3.72% and the other 0.48%; and *PIK3CA, KIT, NRAS*, and *KRAS* mutations were detected in patients with cholangiocarcinoma, hepatocellular carcinoma, gastrointestinal stromal and germ cell tumors, and breast, pancreatic, and head and neck cancers (Supplementary Table S2).

### Association between ctDNA dynamics and clinical outcome

We followed up 106 patients with regards to clinical outcomes. Among them, 26 patients had more than one clinical evaluation, which allowed us to assess the correlation between the changes in ctDNA levels and clinical evaluation at corresponding times. Of the 136 ctDNA datasets, 127 (93.4%) were consistent with the clinical evaluation. The consistency rates reached up to 100% for most of the major tumor types except for intracranial tumors at 71.4% (15/21) and pancreatic cancer at 60.0% (3/5).

For the consistent datasets, although the patients with lung cancer and urinary tract cancer received only two treatments, other patients experienced multiple treatments, including chemotherapy, radiotherapy, chemoradiotherapy, targeted therapy, immunotherapy, and combined chemotherapy and targeted therapy. In addition, some blood samples were taken from patients both just after surgery and during drug withdrawal, defined as postoperation and no treatment, respectively. Most of the surgical patients underwent tumor resection, whereas six received liver transplantation (Table 1). Among these different types of cancers and treatments, the change in ctDNA as determined by the individualized panel was consistent with clinical observations of PD, SD, and remission (Fig. 3A and B). As expected, the consistency rate in the peripheral blood samples was much lower than that in the CSF for glioma patients, at 66.7% and 100%, respectively, reflecting the physical inhibition by the BBB.

**Fig. 3.**
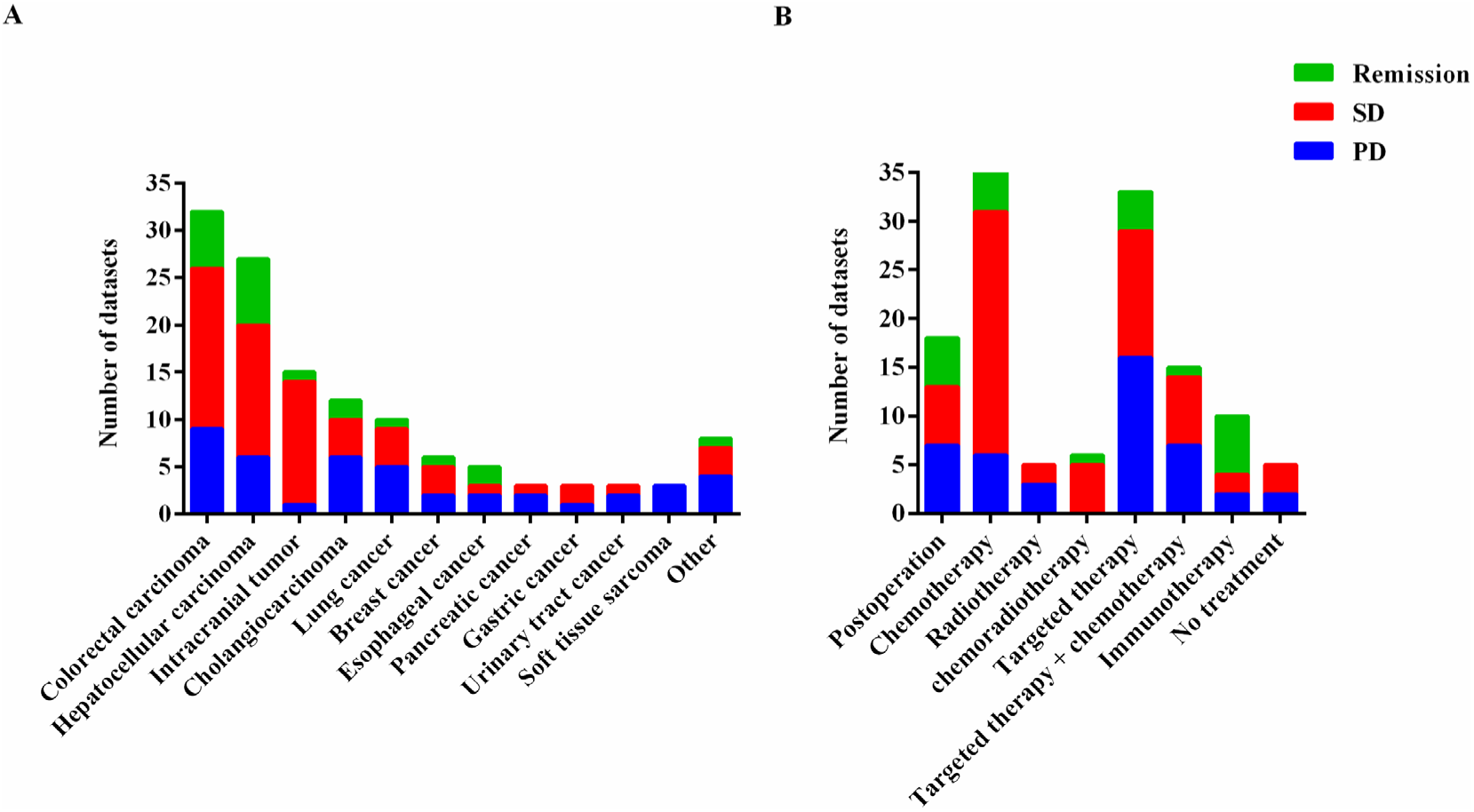
Distribution of treatment responses [progression-free disease (PD), stable disease (SD), and remission] according to different tumor types (A) and treatments (B).

**Table 1.**
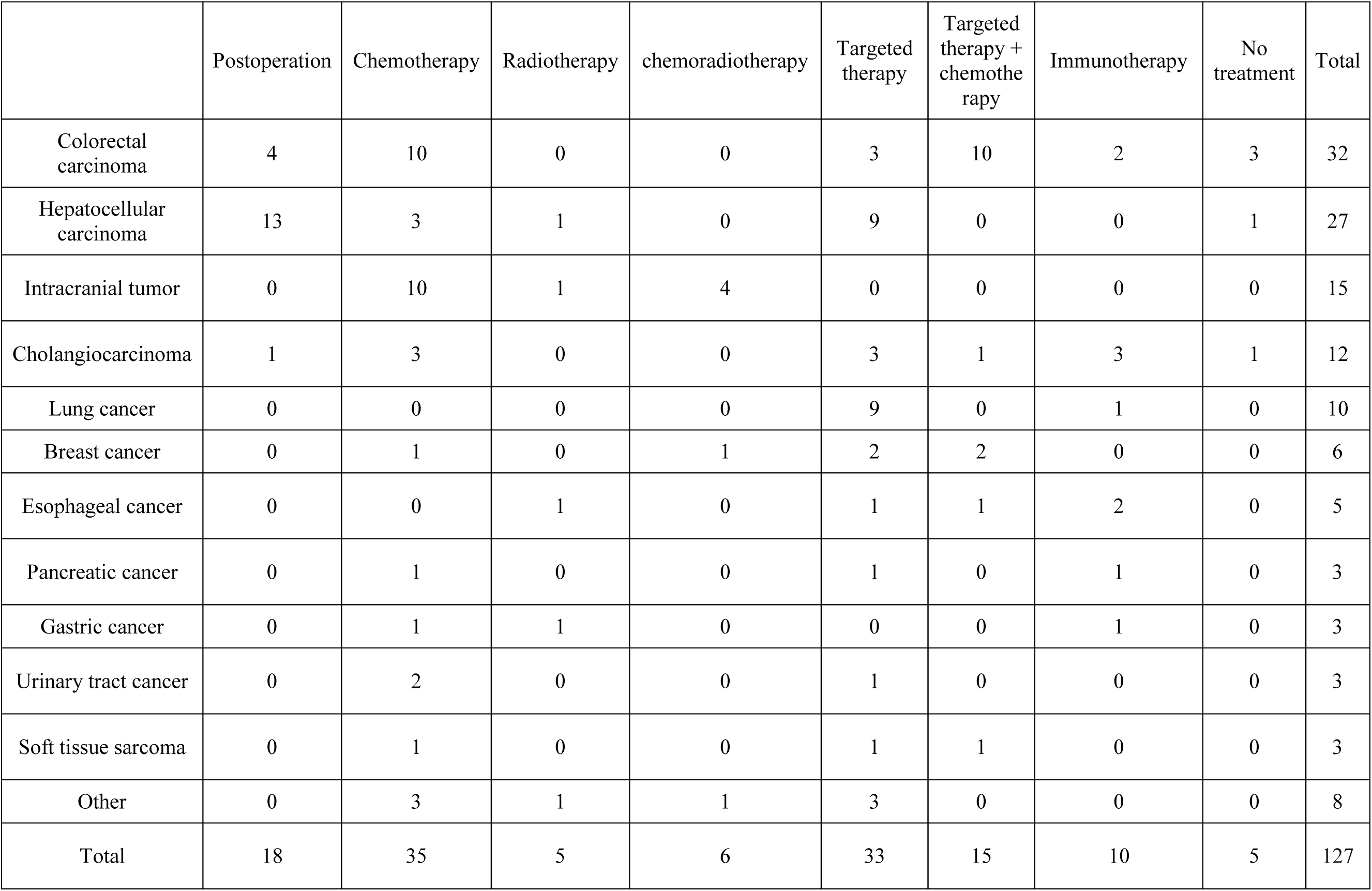
Distribution of treatment types in different cancer types.

We then analyzed the corresponding ctDNA levels with clinical outcome metrics, including PD, SD, and remission. The median CCF of patients with PD was 2.21%. There was no difference in the median CCF between SD patients and those who experienced remission, at 0.17% and 0.31%, respectively (p = 0.144). However, the CCF of the SD and remission patients was one order of magnitude lower than that of the PD patients, representing a statistically significant difference (both p < 0.001) (Fig. 4A).

**Fig. 4.**
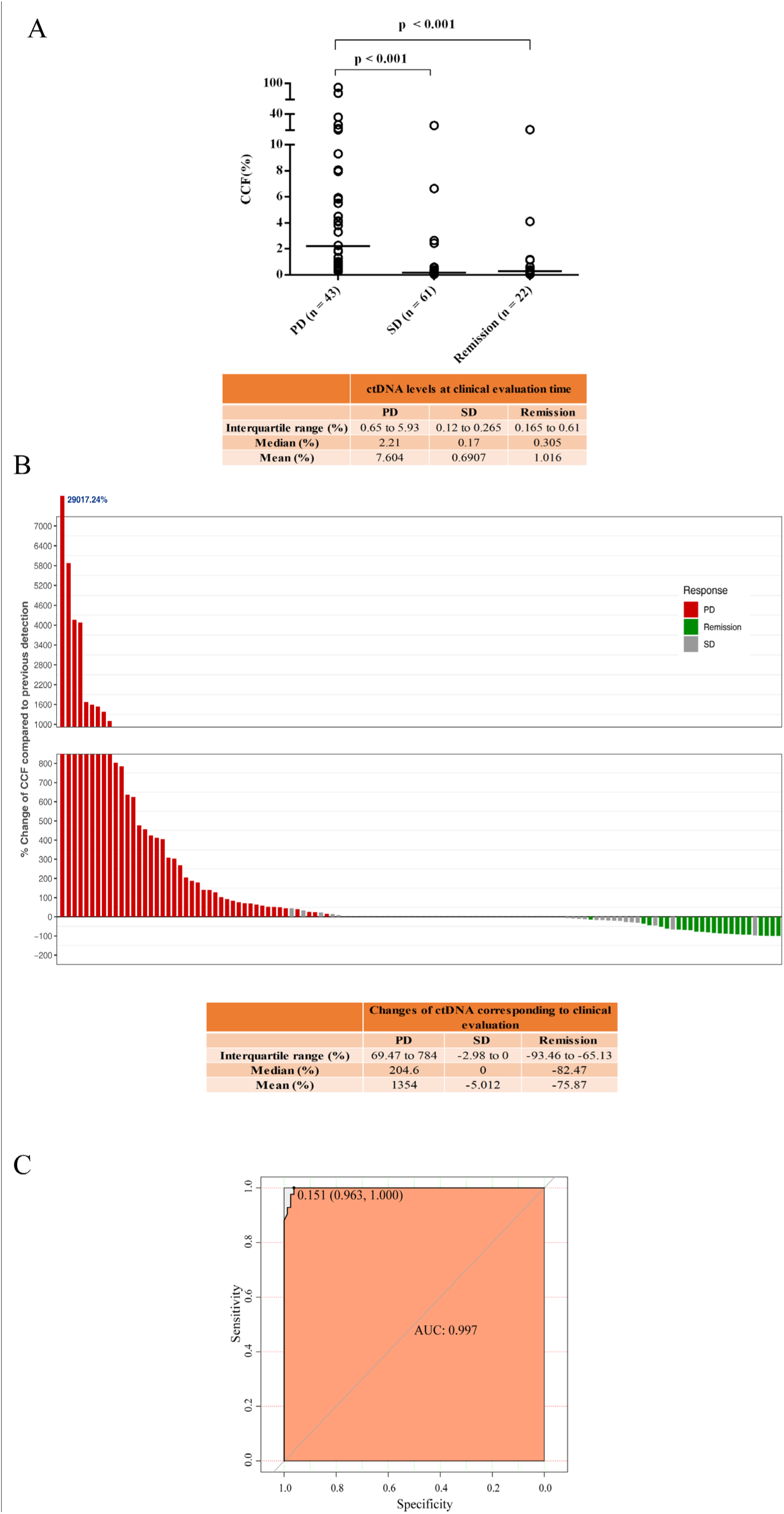
ctDNA concentration change and its use for progression prediction. (A) Distribution of ctDNA concentration in different treatment responses. CCF values were significantly different between PD and SD, and between PD and remission. The p-values were obtained from the Kruskal-Wallis H test. n, number of patients. (B) Change of ctDNA concentration after treatment corresponding to the clinical evaluation. (C) ROC analysis of the ctDNA concentration change. The analysis suggested a threshold value of 15.1% to predict PD.

We also analyzed the percentage change in ctDNA levels between different detection times along with the corresponding time of clinical evaluations. The CCF detected at the time of clinical evaluation significantly differed from the levels detected at the last assessment for the patients with PD and remission, but there was no such difference for the SD patients (p < 0.0001, < 0.0001, and p = 0.0520, respectively; Wilcoxon signed-rank test). Overall, 67.2% of SD patients showed ctDNA levels below the threshold at two time points, which was considered as a change to 0 (Fig. 4B). However, there was high variability among PD patients, with a median change of 204.6% in ctDNA (Fig. 4B). The receiver operating characteristic (ROC) curve suggested that the cut-off value for predicting PD was a 15% increase in the ctDNA level (Fig. 4C). The maximum change of ctDNA observed was a ∼29,000-fold increase, which occurred in a patient with brain metastases of breast cancer (Fig. 4B). After surgical removal of the brain metastases, the patient’s condition was relieved, and the ctDNA level substantially dropped down to 0.29%. However, after 3 months, the ctDNA of this patient again increased to 84.4%, which corresponded with new metastases to the pectoralis major muscle and the brain metastases also recurred.

Among the 43 PD patients, 23 showed metastasis and recurrence (Table 2). In addition to 10 patients who had metastases and relapses identified prior to the first ctDNA test, 13 patients were with metastasis and recurrence were followed-up. Eight patients were at or above the 0.25% threshold at the first ctDNA test, five patients showed a change in the CCF from below the threshold to above and reaching ctDNA-positive status. In summary, all patients with metastasis and recurrence showed a ctDNA-positive status, which could accurately predict metastasis and recurrence. Except for patients P1822 and P2022 who were tested to be ctDNA-positive and underwent imaging metastasis and recurrence at almost the same time, 85% of the patients (11/13) reached the ctDNA-positive status earlier than the imaging diagnosis. The lead time between the first detection of positivity for ctDNA and radiological metastasis or recurrence in patients P2829, P1782, and P1609 was 33, 68, and 128 days, respectively, with an average of 76 days. Moreover, 82.6% (19/23) of the patients had at least one test with a CCF above 1%. Patient P3079 had a ctDNA concentration of 4.99% before the right hepatectomy, and the postoperative ctDNA concentration decreased by 80%, but still considered to be ctDNA-positive, and then the patient experienced relapse after 2 months.

**Table 2.**
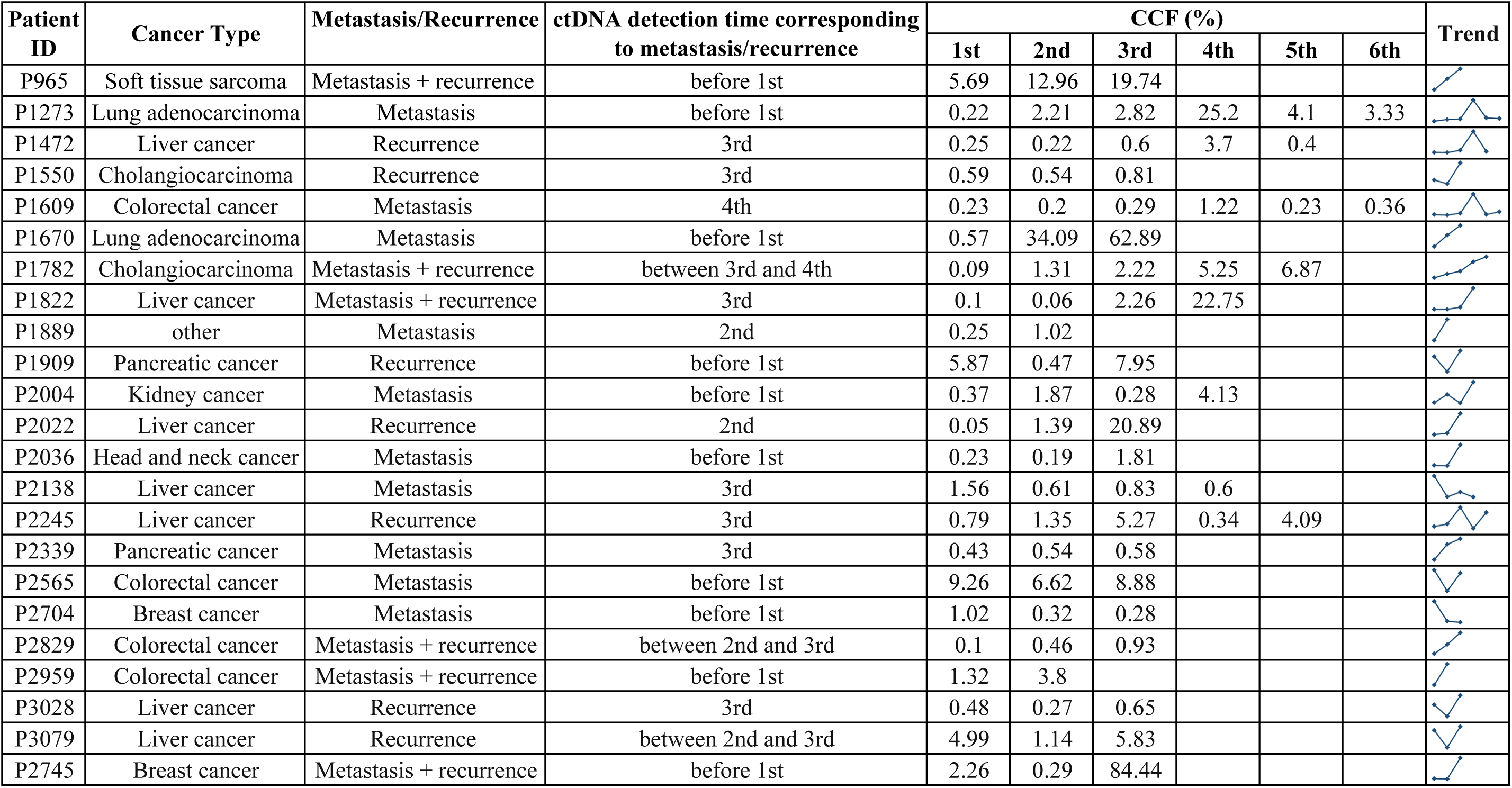
Surveillance of metastasis and recurrence by the individualized ctDNA fingerprint assay.

| Patient ID | Cancer Type | Metastasis/Recurrence | ctDNA detection time corresponding to metastasis/recurrence | CCF (%) |  |  |  |  |  | Trend |
| --- | --- | --- | --- | --- | --- | --- | --- | --- | --- | --- |
|  |  |  |  | 1st | 2nd | 3rd | 4th | 5th | 6th |  |
| P965 | Soft tissue sarcoma | Metastasis + recurrence | before 1st | 5.69 | 12.96 | 19.74 |  |  |  |  |
| P1273 | Lung adenocarcinoma | Metastasis | before 1st | 0.22 | 2.21 | 2.82 | 25.2 | 4.1 | 3.33 |  |
| P1472 | Liver cancer | Recurrence | 3rd | 0.25 | 0.22 | 0.6 | 3.7 | 0.4 |  |  |
| P1550 | Cholangiocarcinoma | Recurrence | 3rd | 0.59 | 0.54 | 0.81 |  |  |  |  |
| P1609 | Colorectal cancer | Metastasis | 4th | 0.23 | 0.2 | 0.29 | 1.22 | 0.23 | 0.36 |  |
| P1670 | Lung adenocarcinoma | Metastasis | before 1st | 0.57 | 34.09 | 62.89 |  |  |  |  |
| P1782 | Cholangiocarcinoma | Metastasis + recurrence | between 3rd and 4th | 0.09 | 1.31 | 2.22 | 5.25 | 6.87 |  |  |
| P1822 | Liver cancer | Metastasis + recurrence | 3rd | 0.1 | 0.06 | 2.26 | 22.75 |  |  |  |
| P1889 | other | Metastasis | 2nd | 0.25 | 1.02 |  |  |  |  |  |
| P1909 | Pancreatic cancer | Recurrence | before 1st | 5.87 | 0.47 | 7.95 |  |  |  |  |
| P2004 | Kidney cancer | Metastasis | before 1st | 0.37 | 1.87 | 0.28 | 4.13 |  |  |  |
| P2022 | Liver cancer | Recurrence | 2nd | 0.05 | 1.39 | 20.89 |  |  |  |  |
| P2036 | Head and neck cancer | Metastasis | before 1st | 0.23 | 0.19 | 1.81 |  |  |  |  |
| P2138 | Liver cancer | Metastasis | 3rd | 1.56 | 0.61 | 0.83 | 0.6 |  |  |  |
| P2245 | Liver cancer | Recurrence | 3rd | 0.79 | 1.35 | 5.27 | 0.34 | 4.09 |  |  |
| P2339 | Pancreatic cancer | Metastasis | 3rd | 0.43 | 0.54 | 0.58 |  |  |  |  |
| P2565 | Colorectal cancer | Metastasis | before 1st | 9.26 | 6.62 | 8.88 |  |  |  |  |
| P2704 | Breast cancer | Metastasis | before 1st | 1.02 | 0.32 | 0.28 |  |  |  |  |
| P2829 | Colorectal cancer | Metastasis + recurrence | between 2nd and 3rd | 0.1 | 0.46 | 0.93 |  |  |  |  |
| P2959 | Colorectal cancer | Metastasis + recurrence | before 1st | 1.32 | 3.8 |  |  |  |  |  |
| P3028 | Liver cancer | Recurrence | 3rd | 0.48 | 0.27 | 0.65 |  |  |  |  |
| P3079 | Liver cancer | Recurrence | between 2nd and 3rd | 4.99 | 1.14 | 5.83 |  |  |  |  |
| P2745 | Breast cancer | Metastasis + recurrence | before 1st | 2.26 | 0.29 | 84.44 |  |  |  |  |

The clinical responses of nine patients were inconsistent with the dynamic changes of ctDNA. These included five patients with gliomas, two with pancreatic cancers, one with medulloblastoma, and one with breast cancer. Although all of the patients with glioma showed disease progression, their plasma ctDNA levels fluctuated below the threshold, owing to the low detection rate of ctDNA in plasma. This suggests that it is necessary to monitor ctDNA in the CSF, instead of the blood, for glioblastoma patients. A similar situation occurred in patients with pancreatic cancer, for whom disease progressed but the CCF was determined to be below the 0.25% threshold. The CCF values of the patient with medulloblastoma changed from 0.19% to 0.34% with an increase rate of 79%. Although this change was close to that observed in PD patients, the status of this patient was SD. For breast cancer, the CCF of the first and second tests was 27.7% and 21.3%, respectively. Although the CCF slightly reduced, the absolute value was still very high, which may reflect the PD status of this patient.

### Dynamics of individual mutations in the individualized ctDNA fingerprint assay

We selected five patients who had at least three clinical evaluations, and analyzed their disease course in detail through tracking their radiological response and ctDNA dynamics, including the overall CCF and individual mutation changes detected in the personalized ctDNA panels. The five patients had lung adenocarcinoma, cholangiocarcinoma with liver transplant surgery, CRC, hepatocellular carcinoma, and breast cancer, respectively, and their changes in ctDNA were consistent with the clinical evaluations (Fig. 5). Patient A received gefitinib for more than one year, and then acquired the EGFR_p.E746_A750del and EGFR_p.T790M mutations. Before imaging confirmed PD in May of 2018, the fraction EGFR_p.T790M mutation slightly increased and then decreased, whereas the EGFR_p.E746_A750del mutation fraction increased continuously. After treatment with osimertinib, the patient achieved a CR and maintained low ctDNA levels. Moreover, the fluctuations in certain mutations observed in patient D and patient E differed from the overall trend of the panel. For patient D, the ctDNA levels of *DPYD* and *IGSF1* increased significantly in the second detection, and that of *PCSK5* increased in the third test, but the overall ctDNA content of the panel decreased in the second and third detections. Patient E also contained an *AP1M2* mutation that was distinct from the overall trend of ctDNA changes.

**Fig. 5.**
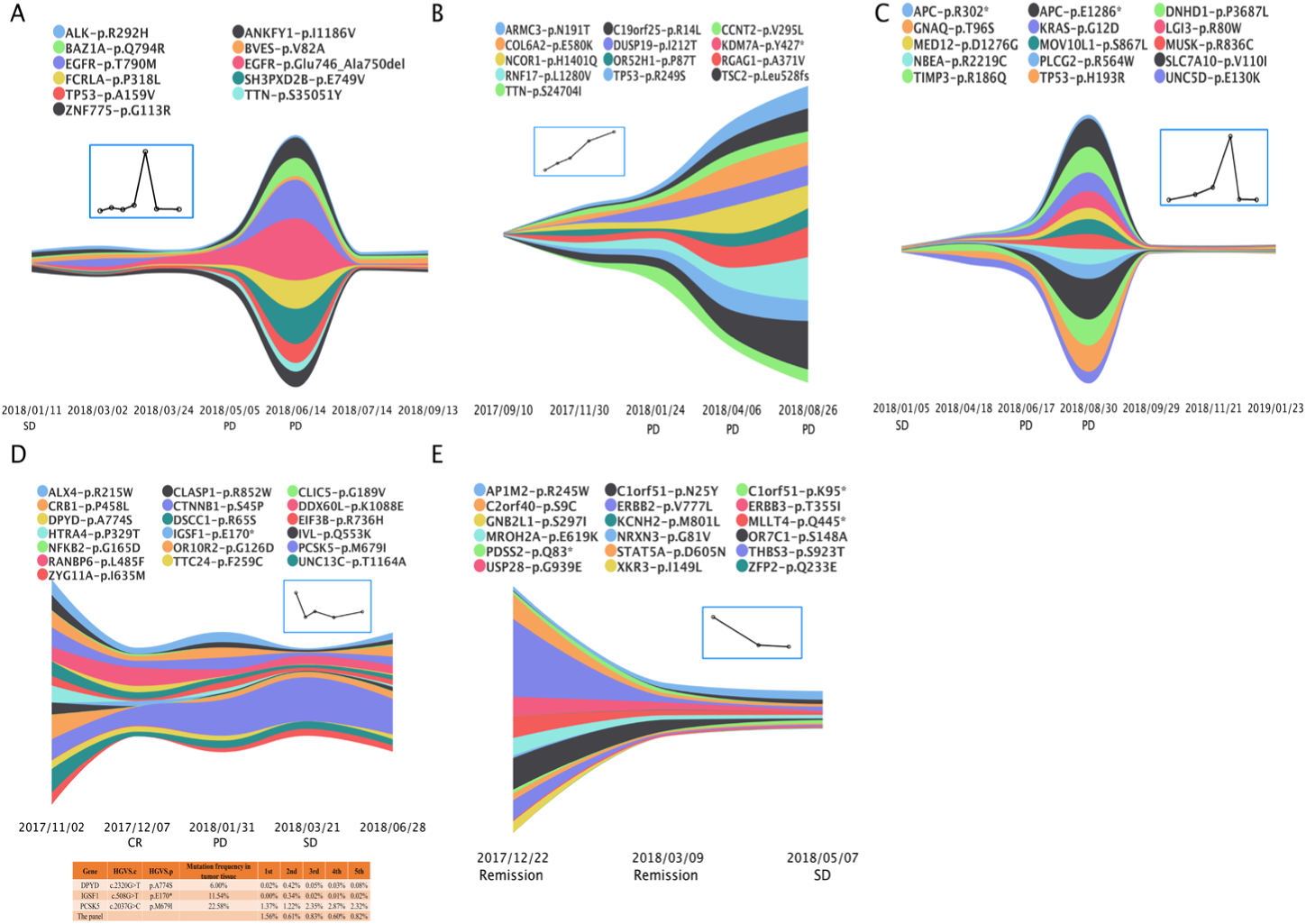
Stream graph of five patients showing dynamics of individual mutations as well as the overall trend (inset line plot) as detected by ctDNA. The colors indicate different tumor genes and the specific mutations. (A) Lung adenocarcinoma; (B) cholangiocarcinoma with liver transplant surgery; (C) CRC; (D) hepatocellular carcinoma; and (E) breast cancer. For patient D, a table presents the dynamics of three mutations, which were inconsistent with the overall trend.

### Patient individualization of customized ctDNA panels

We designed the individualized ctDNA panels based on the patients’ own WES data, and analyzed the specificity of this individualization. A total of 144,158 mutations were detected in 521 patients by WES, including 99,490 mutations that were detected only once, accounting for 69.0% of all mutations. The 99,490 unique mutations were successfully incorporated in at least one individualized ctDNA panel, and 7762 (84.9%) of the mutations were incorporated in only one panel; thus, these panels were highly specific to the individual patients.

To calculate the occurrence of common tumor-associated genes in the individualized ctDNA panel, we defined a tumor-associated gene set using the following method. Based on a previous survey including approximately 90 commercially available clinical gene panels developed by 40 different research institutes and companies, a list was generated containing 2054 genes (11). Using this list, we subsequently screened for 377 genes that occurred five or more times in all commercial panels (Supplementary Table S3). These genes were considered to be tumor-associated genes and potentially clinically actionable. Mutations with frequencies greater than 5% were selected from the WES data and compared to these 377 genes; 114 (21.8%) patients did not contain any of the 377 gene mutations on the list, and 365 (70%) patients had 1– 3 of these genes, while 42 (8%) patients had more than three of the genes on the list. Similar results were obtained when comparing these 377 genes with the genes in the individualized ctDNA panels.

## Discussion

Although proof-of-concept studies have confirmed the clinical value of ctDNA to predict the treatment response (12), an expert panel from the American Society of Clinical Oncology and the College of American Pathologists concluded that most of the clinical validity and practicability of ctDNA testing reported to date are insufficient (13). Several challenges have to be overcome before ctDNA analysis can be translated to clinical application. Toward this end, we developed a strategy to design patient-specific ctDNA panels with high sensitivity and specificity, which can be applied to patients with multiple types of tumors that received several different treatments.

Our panels achieved a specificity of 99.99% when using 10 SNVs with a detection frequency above 0.25%. The specificity was 99.93% in the reference samples. A median of 19 SNVs were detected in our ctDNA fingerprint assays. The ctDNA detection rate reached up to 67% in all patients, and about 80% in patients with CRC, non-small cell lung cancer (NSCLC), and gastric cancer. In comparison, a targeted deep-sequencing study of more than 70 tumor-associated genes in more than 20,000 patients with solid tumors showed a ctDNA detection rate of 85% overall, and 90% in patients with liver cancer, NSCLC, CRC, and gastric cancer (14). The higher detection rate in that study can be attributed to the fact that all of the included patients had been treated for late-stage cancers. This previous study also showed that the ctDNA detection rate was relatively low in brain cancer patients due to the BBB (14), which is consistent with our present findings.

Previous studies showed that ctDNA could indicate the treatment response and detect recurrence earlier than radiographic imaging in CRC patients that received surgery or chemotherapy (15–17). In a study of 20 patients with metastatic breast cancer, 95% of the patients showed ctDNA fluctuations consistent with imaging-determined PD and SD using a patient-specific ctDNA panel, and 53% of the patients showed an increase in ctDNA that occurred earlier than the imaging-detected PD (18). Monitoring with a panel of 61 hotspot genes revealed that a ctDNA increase was associated with clinical relapse and PD in patients with muscle-invasive bladder cancer (19). These studies generally focused on only one type of cancer or one treatment. In contrast, in the present study, we collected data from patients with more than 12 tumor types and performed ctDNA monitoring on seven different treatment options, including liver transplantation. The ctDNA levels we detected in patients with PD, remission, and SD status were significantly different from each other. At the same time, the correlation between the changes of ctDNA level and clinical evaluation was high at 93%.

Immunotherapy has shown great value in cancer treatment and has been approved by the Food and Drug Administration of the USA for the treatment of a variety of cancers (20, 21). Since it is expected to be applied to a large and growing number of patients, there is a large demand to develop effective methods for monitoring the efficacy of immunotherapy. Monitoring ctDNA changes is a promising approach to begin filling this gap. In a recent study in advanced NSCLC patients treated with nivolumab, 100% of the patients showing an objective response (OR) to immunotherapy had reduced ctDNA concentrations with a median change of 87.8%, while 60% of the non-OR patients had increased ctDNA concentrations (22). In the present study, our individualized ctDNA panels successfully tracked PD, remission, and SD in patients with digestive tract cancers who received treatment with PD-L1 blockers and cell immunotherapy. The consistency rate reached up to 100% (10/10) between the trends of ctDNA and disease status change (Fig. 3). Thus, this method can effectively monitor multiple cancer types and treatment modes.

The ctDNA panels also have advantages over monitoring only single mutations in predicting disease development. The dynamics of a single mutation representation can be inconsistent with clinical assessments in such cases as well as the overall trend of multiple mutations. In a breast cancer patient whose status changed from SD to PD, the ctDNA content of most mutations increased, but the level of *ZFYVE21*, a known breast cancer marker, continued to decrease (18). Similarly, the *NRAS* level, a known melanoma marker, was reduced by 10-fold in a melanoma patient, but the patient’s disease nevertheless progressed (23). These reports are consistent with our findings (Fig. 5D-E). Hence, we monitored multiple patient-specific mutations to avoid this problem.

In addition to monitoring disease status, our ctDNA panels also have potential to predict patient prognosis at an earlier stage than currently possible. The ROC analysis identified a 15.1% increase in ctDNA as the optimal threshold for predicting PD. The corresponding AUC value was 0.997, suggesting a high confidence level (Fig. 4C). Importantly, except for patients with recurrence or metastasis identified before the first ctDNA test, 85% of the patients (11/13) with a ctDNA-positive status at the first ctDNA test or who converted to a ctDNA-positive status during follow-up experienced recurrence or metastasis far before these events were detected by imaging. The remaining patients tested as ctDNA-positive and demonstrated recurrence and metastasis by imaging at almost the same time. The average lead time from the first ctDNA-positive finding to radioactive-confirmed metastasis and recurrence in three patients who reached a ctDNA-positive status during follow-up was 76 days. This finding is similar to a previous study in which 68 patients with advanced bladder cancer were followed-up and ctDNA monitoring showed a median lead time of 107 days (24). Given the relatively small sample size of this study, the clinical advantage of individualized ctDNA panels in predicting metastasis and recurrence need to be further explored on a larger scale in the future.

The cost of using individualized ctDNA panels is expected to be much lower than several previously reported ctDNA panels such as CAncer Personalized Profiling by deep Sequencing (CAPP-Seq), targeted error correction sequencing (TEC-Seq), and a commercially developed ctDNA test kit including more than 50 genes (25–28). All of these panels were designed to target tumor hotspot genes, which account for only a minority of the total ctDNA in the plasma. In cancer patients, somatic cell-free DNA (cfDNA) consists of variants derived from both clonal hematopoiesis and the tumor. Although the common tumor mutations also occur in the genome of white blood cells at high frequency, the majority of cfDNA variants are nevertheless derived from white cells produced in clonal hematopoiesis (29, 30). Therefore, detection based on common tumor mutations is of low efficiency and high cost.

In our present study, we used paired WES data from patients to select variants free of clonal hematopoietic cell interference. The resulting ctDNA panels customized for individual patients displayed high specificity for all patients; 85% of the selected mutations were detected only once, about 30% of the patients did not harbor common hotspot tumor genes, and the remaining 70% patients carried only 1–3 hotspot genes. This detection efficiency was achieved by sequencing a total of about 30 patient-specific mutations per individual, and each of these mutations was harbored in a 100-bp amplicon. Thus, the overall sequencing and analysis involves only about 3-kb sequences per individual. This greatly reduces the cost and time of analysis in comparison to those required for panels needing larger sequencing volumes (31).

Nevertheless, the present individualized ctDNA fingerprints assay still has some limitations. Importantly, it requires prior WES analysis of the tumor tissue before ctDNA monitoring, which is impossible for some patients. Additionally, this assay only uses information of SNVs due to the short sequencing length and leaves out information about indels and longer range changes, including gene fusions/chromosomal rearrangements and copy number variations (CNVs), which are highly valuable diagnosis tools for some tumors (32, 33). Although the present platform mainly detects SNVs, there is relatively less information available on related fusions and CNVs. Furthermore, the present study was limited to patients with metastasis and recurrence, which requires further exploration of the full breadth of clinical benefits of these individualized ctDNA panels.

In conclusion, we developed a new ctDNA platform that could successfully monitor the treatment response, recurrence, and metastasis in a variety of cancer types and for patients receiving multiple treatments. Although further large-scale clinical trials are needed to confirm the utility of these individualized ctDNA panels in clinical settings, the panels offer the important advantages of convenience, accuracy, and efficiency, and show promise for improving personalized treatment decisions in a timely manner.

## Material and Methods

### Patient samples

Patients were recruited from 20 hospitals in China between May 2016 and December 2018. All patients provided written informed consent to use their genomic and clinical data for research purposes. All procedures involving human participation conformed to the ethical standards of the relevant institutions and/or National Research committees, as well as the 1964 Helsinki Declaration and its subsequent amendments or similar ethical standards.

Tumor tissues and blood samples of 552 Chinese patients diagnosed with cancer were collected. The percentage of tumor cells was assessed by pathologists, and only samples with a tumor content greater than 20% were included in this study. A series of plasma samples were collected at intervals set by the treating physicians. Treatment response and recurrence were evaluated by oncologists based on radiological imaging. ctDNA tests were conducted within the same month of radiological imaging for over 90% of the cases. In the remaining 10% of cases, the time between the ctDNA test and radiological imaging was between 1–2 months. All patients were followed up for a median of 6 months (range 1–16 months).

Reference samples used to evaluate the ctDNA detection threshold and performance were prepared from commercial references including Quantitative Multiplex Formalin Compromised (Mild) Reference Standard (cat# HD798, Horizon Discovery, Cambridge, UK), 1% Multiplex I cfDNA Reference Standard (cat# HD778, Horizon Discovery, Cambridge, UK), and 100% Wildtype (Tru-Q 0) (cat# HD752, Horizon Discovery, Cambridge, UK). The reference standards were mixed in different proportions to make serial reference samples including different variants with a gradient allelic frequency of 15%, 10%, 6%, 3%, 1%, 0.5%, 0.25%, and 0%, respectively. Variant frequencies detected in ctDNA measurements were then used to calculate the sensitivity and specificity of the assay.

### DNA extraction

For WES analysis, DNA was extracted from the tumor tissue and blood samples previously frozen using Maxwell® RSC Tissue DNA Kit (cat# AS1610, Promega, Madison, WI, USA) and Maxwell® RSC Blood DNA Kit (cat# AS1400, Promega, Madison, WI, USA) respectively. In addition to fresh tumor tissues, the samples included formalin-fixed, paraffin-embedded (FFPE) tissues, which can damage the quality of DNA; therefore, we used NEBNext FFPE DNA Repair Mix (cat# M6630, New England Biolabs, Ipswich, MA, USA) to reduce this damage. For the ctDNA assay, 10 ml of whole blood was collected and transported using Streck’s BCT tubes. cfDNA in the plasma and CSF was then extracted using MagMAX™ Cell-Free DNA Isolation Kit (cat# 29319, ThermoFisher Scientific, Waltham, MA, USA) and stored at −20°C until use.

### WES analysis

The whole exons of the tumor tissue and matched white blood cells were sequenced. Library preparation was performed using SureSelect Human All Exon Kit v5 (cat# 5990-9857EN, Agilent Technologies, Santa Clara, CA, USA), sequencing was performed on the Illumina Xten platform, and data analysis was carried out as described previously (34).

### ctDNA panels

Given the limitations of multiplex PCR platforms for producing homopolymer-associated indel errors (35, 36), the individualized ctDNA panel was mainly designed to detect somatic SNVs in the peripheral blood. We calculated the clonal clusters for each somatic mutation obtained from WES using SciClone tools, which computationally determines the number and composition of genetic clones across one or multiple samples. The clonal clusters were calculated with the following parameters: tumor purity reviewed by pathologists, tumor copy number alterations (CNA), loss of heterozygosity (LOH) ratio in somatic mutation regions, and somatic variant allele frequencies (VAF). Regions of CNA and LOH were inferred from WES data for each patient using VarScan v2.4.2 tools (37). We selected the top 20–40 somatic mutations within high clonal population clusters, which were applied to Ion AmpliSeq™ Designer for multiplex PCR and amplicon sequencing.

The pre-designed ctDNA panel contains tumor hotspot markers *BRAF, EGFR, ERBB2, KIT, KRAS, MET, NRAS*, and *PIK3CA* to monitor acquired resistance mutations. The information of specific genetic mutations is provided in Supplementary Table S4.

### Multiplex PCR and amplicon sequencing

A total of 5–10 ng of cfDNA was used as the template in the multiplex PCR and amplified using KAPA2G Fast Multiplex Mix kit (cat# KK5802, KapaBiosystems, Wilmington, MA, USA) for the individualized ctDNA panel and Ion AmpliSeq™ Library Kit 2.0 (cat# 4475345, ThermoFisher Scientific, Waltham, MA, USA) for the pre-designed panel. Amplified products were ligated to barcoded adapters from Ion Xpress Barcode Adapters 1-16 Kit (cat# 4471250, ThermoFisher Scientific), followed by NGS library amplification using KAPA HiFi HotStart ReadyMixPCR Kit (cat# KK2602, KapaBiosystems) for pre-designed ctDNA panel and Ion AmpliSeq™ Library Kit for individualized ctDNA, respectively. All libraries were sequenced using Ion Torrent next-generation sequencing platforms (Ion S5 Systems, ThermoFisher Scientific).

### CCF estimation

After sequencing, high-quality data were retained for analysis according to the following criteria: (1) more than 70% of bases scoring Q20 and above; (2) more than 40% of reads mapped to the designed amplicons; (3) average coverage of amplicons higher than 18,000×; (4) >80% amplicons with a coverage above 1000×. The human reference genome hg19 (Feb.2009 GRCh37/hg19) was used for reads alignment with BWA-0.7.12 (38). Samtools-0.1.18 (39) and bedtools-2.17.0 (40) were used to manipulate alignments, and SNVs were called with VarScan v2.4.2 (37) using the mpileup2snp program. We estimated the ctDNA content fraction in patients according to the following formula: 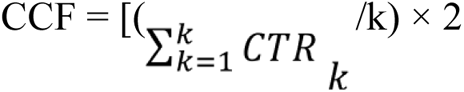, in which CTR refers to the selected somatic mutation allele ratio in the ctDNA test, and k refers to the number of mutations.

## Supplementary Materials

**Supplementary Table S1.** Analytical performance and threshold of the multiplex PCR ctDNA assay for single mutation detection.

| **ctDNA concentration of standard** | 0% | 0.10% | 0.25% | 0.50% | 1% | 3% | 6% | 10% | 15% |
| --- | --- | --- | --- | --- | --- | --- | --- | --- | --- |
| **Number of replicates** | 157 | 160 | 160 | 160 | 157 | 155 | 157 | 157 | 157 |
| **Observed allelic frequency** |  |  |  |  |  |  |  |  |  |
| Mean | 0.06465 | 0.1221 | 0.2761 | 0.5774 | 1.132 | 3.38 | 6.358 | 9.858 | 15.26 |
| Std. Deviation | 0.06247 | 0.1126 | 0.1679 | 0.2678 | 0.1966 | 0.4661 | 0.7514 | 0.9003 | 1.142 |
| Std. Error of Mean | 0.004985 | 0.008899 | 0.01327 | 0.02117 | 0.01569 | 0.03744 | 0.05997 | 0.07186 | 0.09117 |
| Lower 95% CI of mean | 0.0548 | 0.1045 | 0.2499 | 0.5356 | 1.101 | 3.306 | 6.239 | 9.716 | 15.08 |
| Upper 95% CI of mean | 0.0745 | 0.1397 | 0.3023 | 0.6192 | 1.163 | 3.454 | 6.476 | 10 | 15.44 |
| Minimum | 0 | 0 | 0 | 0.02 | 0.66 | 2.1 | 4.74 | 7.66 | 11.69 |
| 25% Percentile | 0.01 | 0.03 | 0.1225 | 0.3725 | 1.01 | 3.05 | 5.8 | 9.29 | 14.31 |
| Median | 0.05 | 0.07 | 0.285 | 0.555 | 1.11 | 3.41 | 6.31 | 9.76 | 15.26 |
| 75% Percentile | 0.105 | 0.2075 | 0.4 | 0.76 | 1.255 | 3.67 | 6.95 | 10.38 | 15.93 |
| Maximum | 0.23 | 0.41 | 0.76 | 1.43 | 1.77 | 4.94 | 8.2 | 14.43 | 18.49 |
| **Coefficient of variation** | 96.62% | 92.17% | 60.81% | 46.39% | 17.37% | 13.79% | 11.82% | 9.13% | 7.49% |
| **Sensitivity (% , threshold**<br>**= 0.127%)** | NA | 40.62% | 75.00% | 96.25% | 100% | 100% | 100% | 100% | 100% |

**NA = not applicable**

**Supplementary Table S2.** Acquired drug-related mutations in 16 patients using a pre-designed ctDNA panel

| Patient ID | Tumor Type | New mutation | Mutation frequency |
| --- | --- | --- | --- |
| tdP1499 | Colorectal cancer | *KRAS* c.183A>C p.Q61H | 0.86% |
| P2074 | Colorectal cancer | *KRAS* c.34G>T p.G12C | 0.60% |
| P2230 | Colorectal cancer | *KRAS* c.34G>T p.G12C | 0.43% |
| P2800 | Colorectal cancer | *KRAS* c.38G>A p.G13D | 0.68% |
| P2997 | Colorectal cancer | *KRAS* c.37G>T p.G13C | 1.75% |
| P3439 | Colorectal cancer | *KRAS* c.183A>T p.Q61H | 1.40% |
|  |  | *NRAS* c.182A>T p.Q61L | 0.98% |
|  |  | *BRAF* c.G1780A p.D594N | 0.69% |
| P3489 | Lung adenocarcinoma | *EGFR* c.2369C>T p.T790M | 3.72% |
| P0220 | Lung adenocarcinoma | *EGFR* c.2369C>T p.T790M | 0.48% |
| P3260 | Lung adenocarcinoma | *PIK3CA* c.1636C>A p.Q546K | 2.41% |
| P2842 | Breast cancer | *PIK3CA* c.G1624A p.E542K | 12.35% |
| P3416 | Pancreatic cancer | *KRAS* c.35G>A p.G12D | 3.46% |
|  |  | *NRAS* c.181C>A p.Q61K | 0.99% |
|  |  | *NRAS* c.182A>T p.Q61L | 0.95% |
| P3490 | Head and neck cancer | *PIK3CA* c.3139C>T p.H1047Y | 0.51% |
| P3954 | Gastrointestinal stromal tumor | *KIT* c.2467T>A p.Y823N | 2.56% |
| P4789 | Cholangiocarcinoma | *PIK3CA* c.1634A>G p.E545G | 0.82% |
| P5513 | Germ cell tumor | *KIT* c.2466T>G p.N822K | 3.81% |
| P2393 | Hepatocellular carcinoma | *NRAS* c.35G>A p.G12D | 0.32% |

**Supplementary Table S3.**
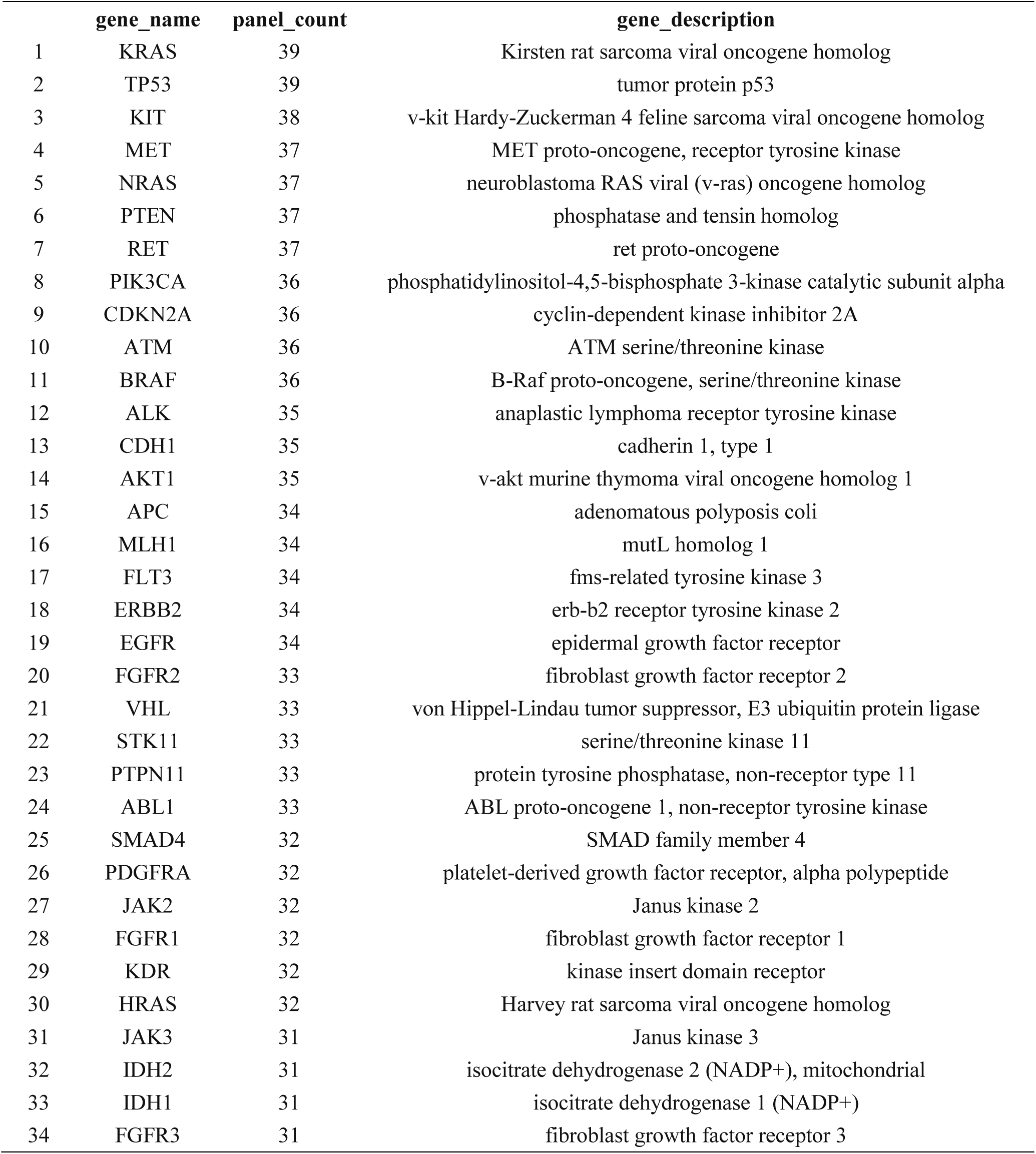
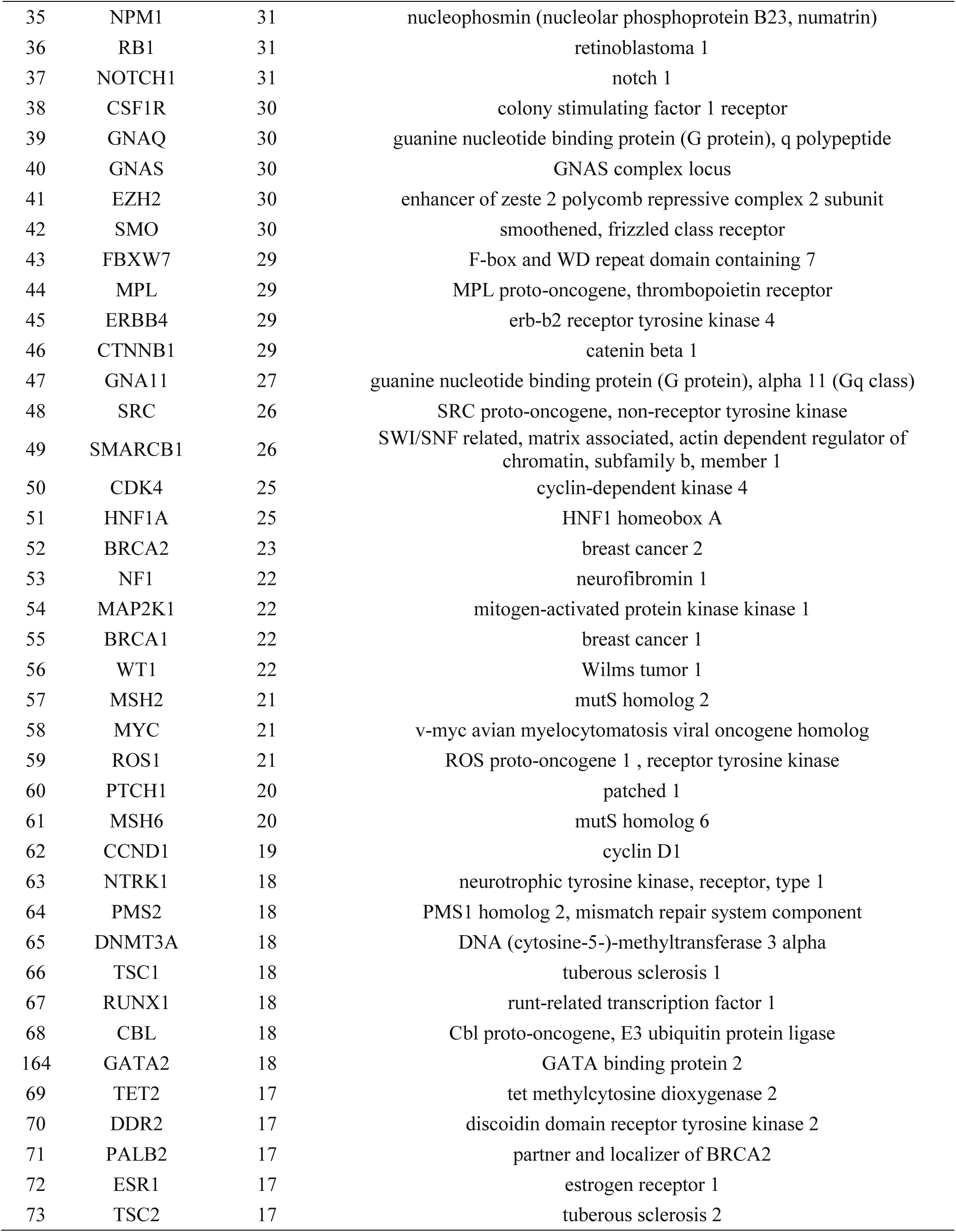
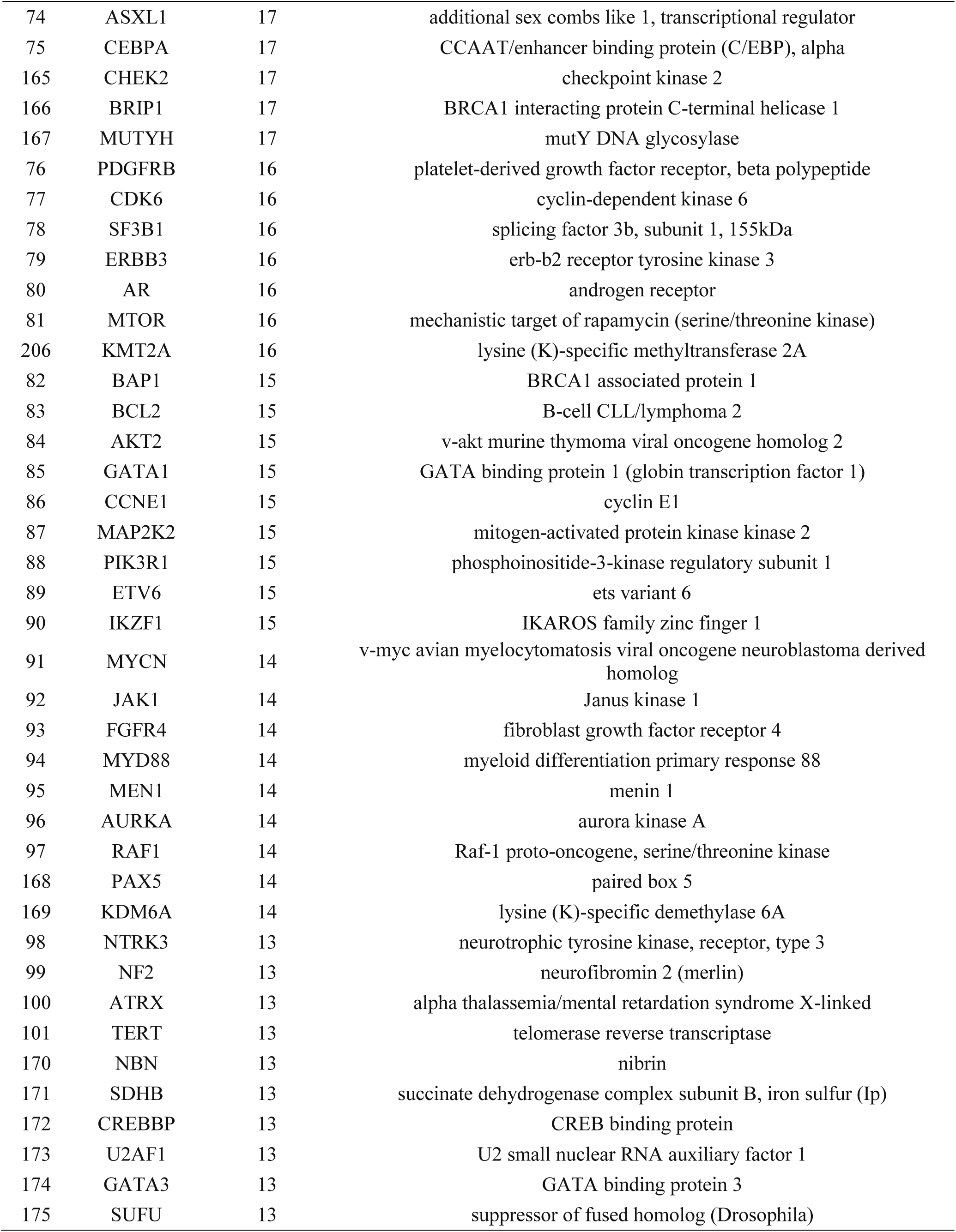
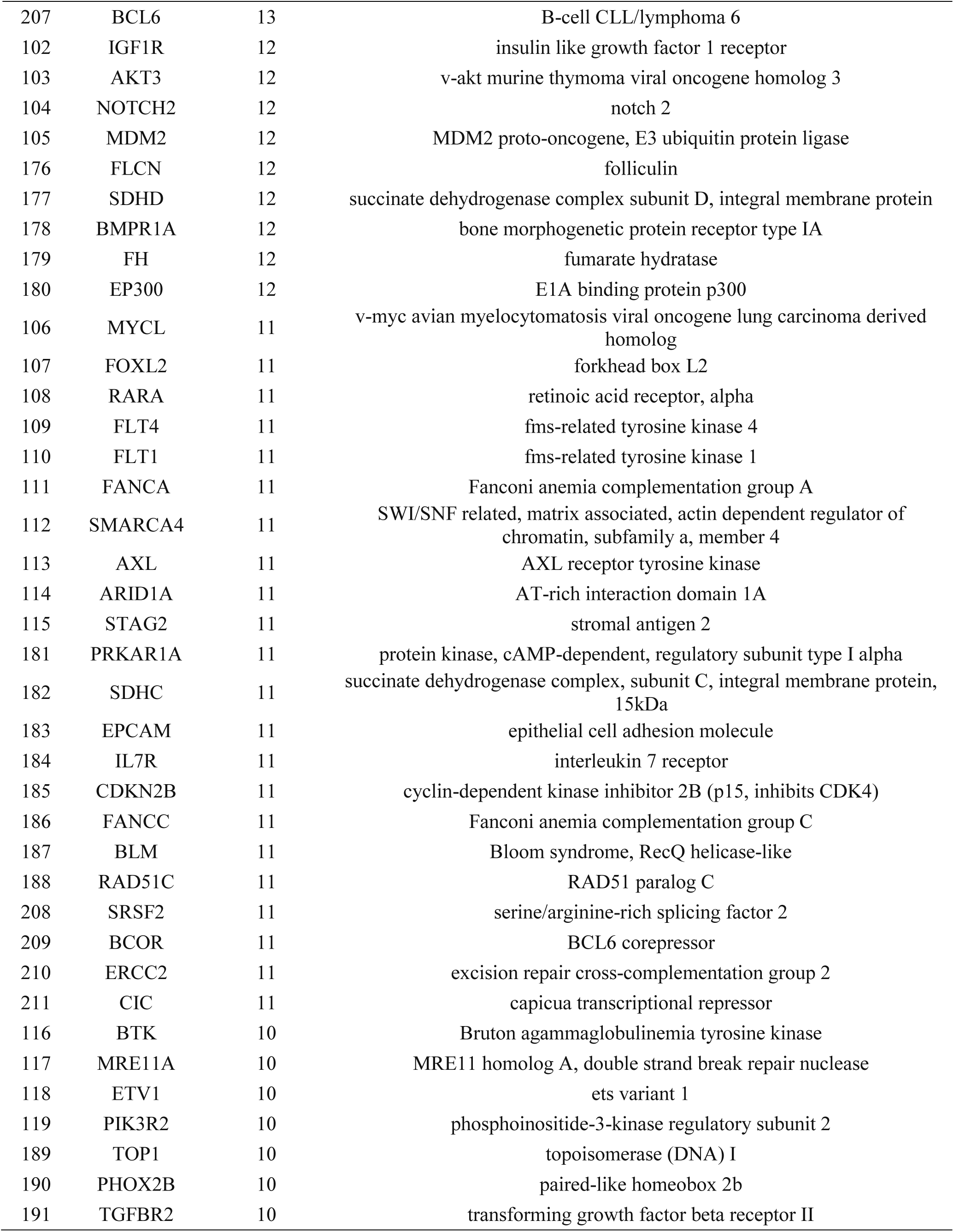
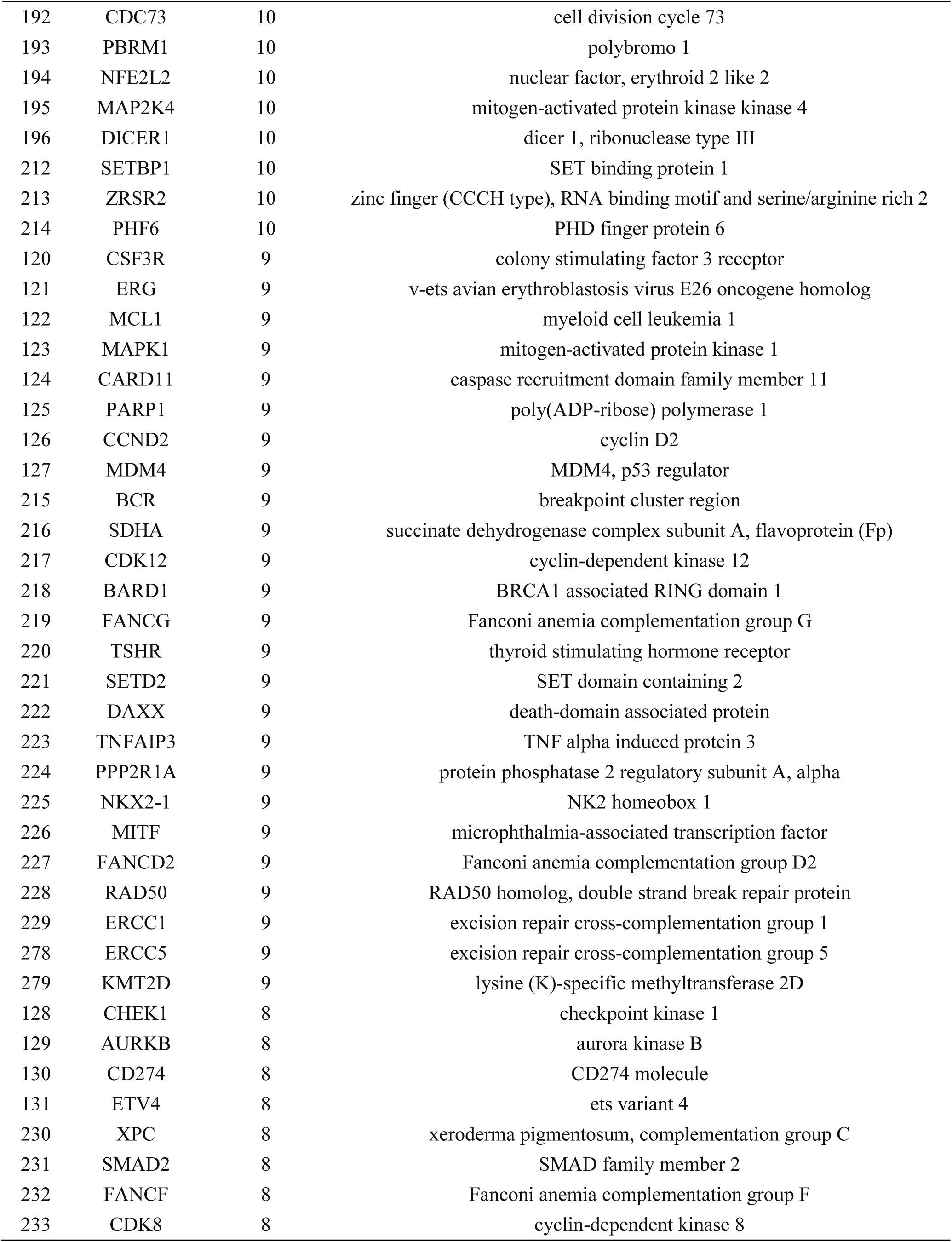
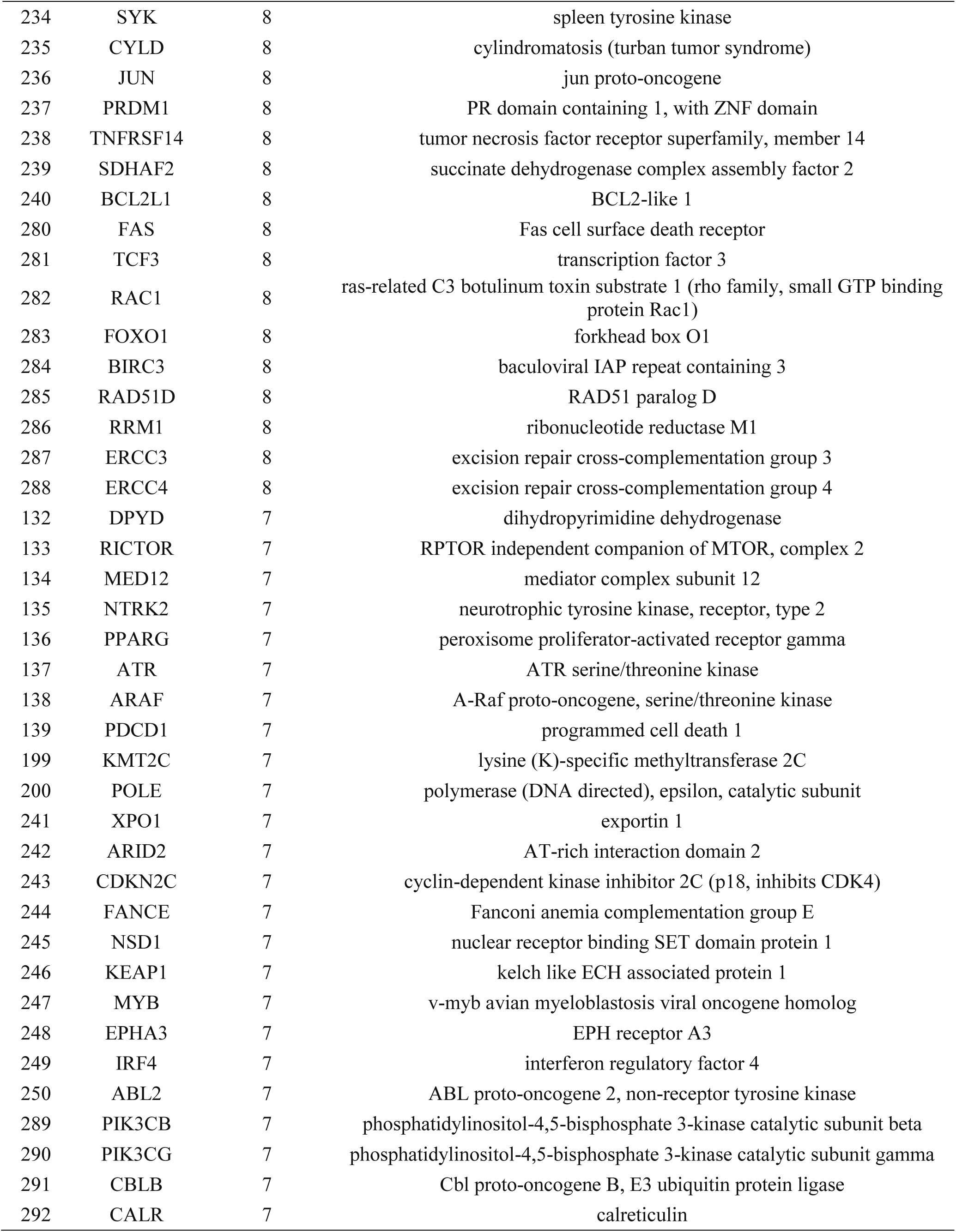
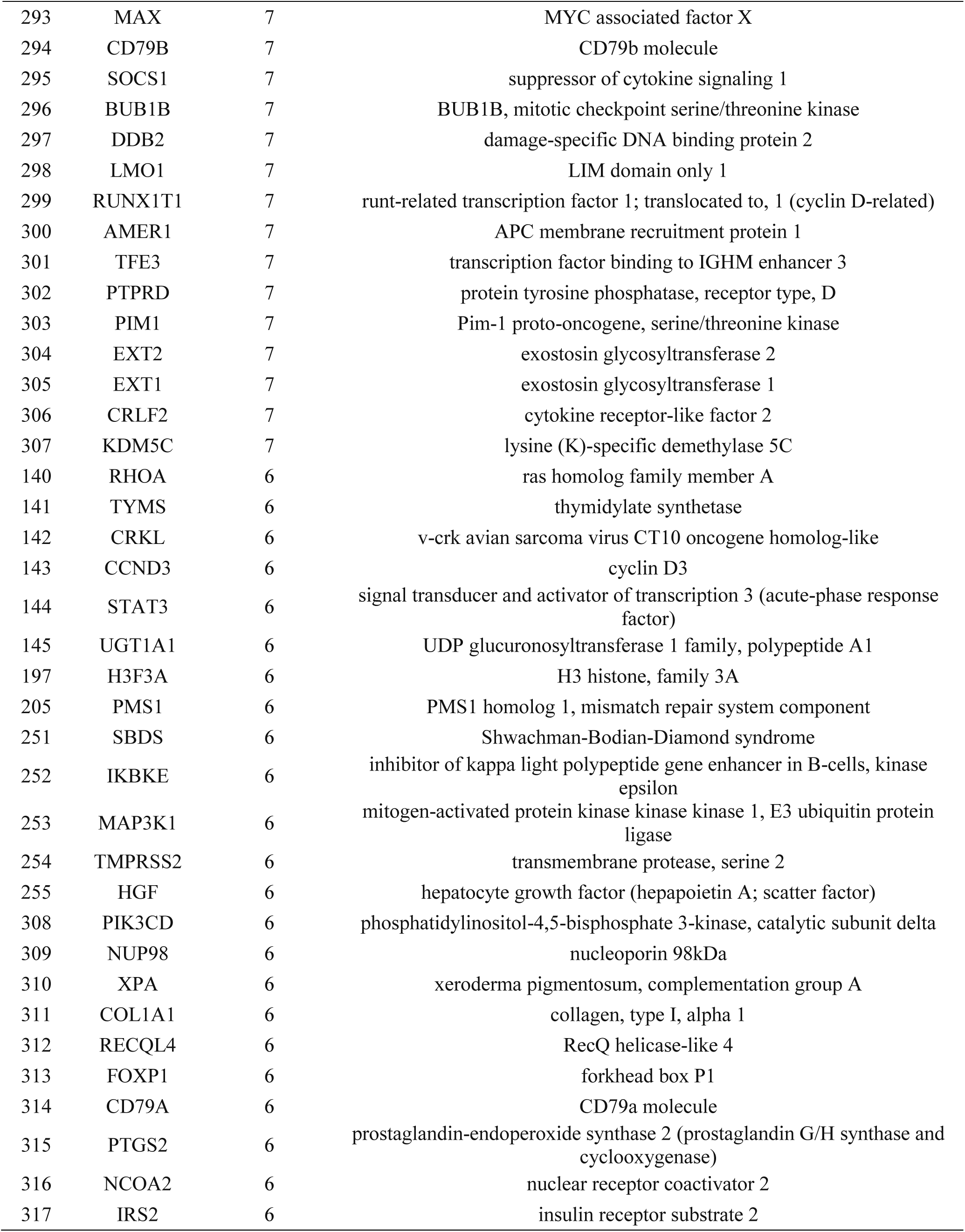
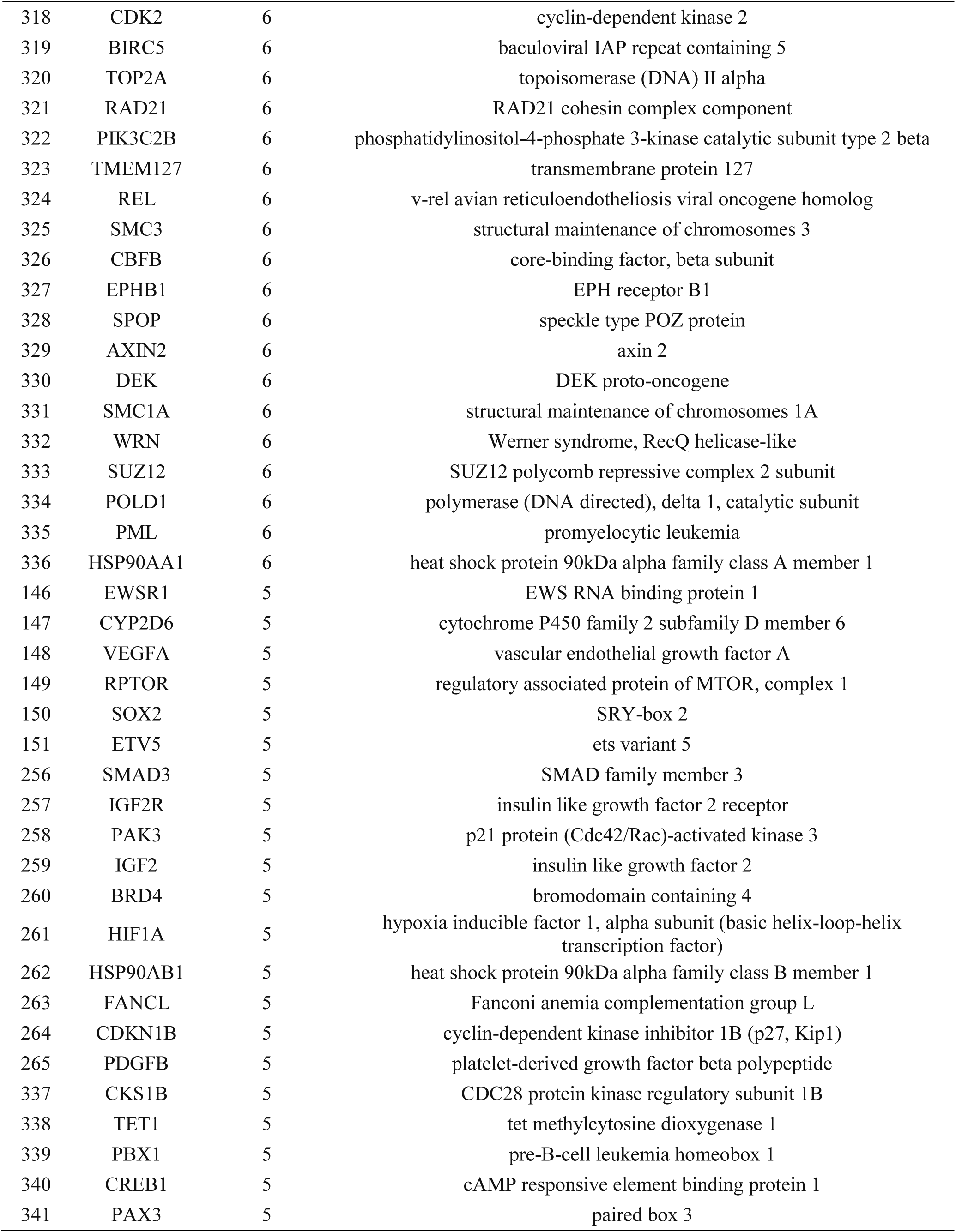
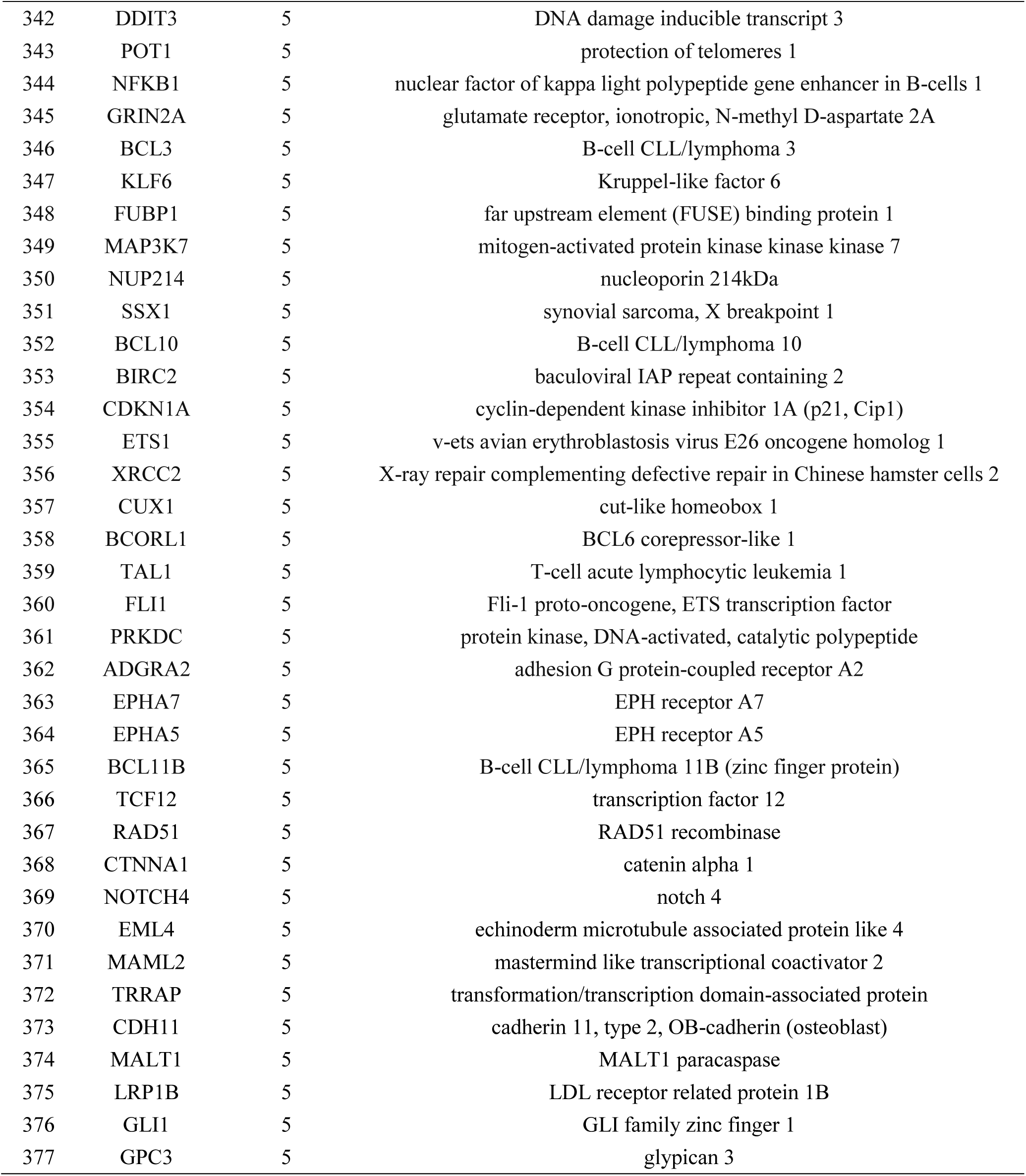
A total of 377 genes that occurred five times or more in a survey of 90 commercially available clinical gene panels developed by 40 different research institutes and companies.

**Supplementary Table S4.** Mutations in the pre-designed ctDNA panel

| gene | CDS Mutation | AA Mutation | chrom | start | end |
| --- | --- | --- | --- | --- | --- |
| BRAF | c.1803A>T | p.K601N | 7 | 140453132 | 140453132 |
| BRAF | c.1801A>G | p.K601E | 7 | 140453134 | 140453134 |
| BRAF | c.1799_1801delTGA | p.V600_K601>E | 7 | 140453134 | 140453136 |
| BRAF | c.1799_1800TG>AA | p.V600E | 7 | 140453135 | 140453136 |
| BRAF | c.1799_1800TG>AT | p.V600D | 7 | 140453135 | 140453136 |
| BRAF | c.1799T>A | p.V600E | 7 | 140453136 | 140453136 |
| BRAF | c.1798_1799GT>AA | p.V600K | 7 | 140453136 | 140453137 |
| BRAF | c.1798_1799GT>AG | p.V600R | 7 | 140453136 | 140453137 |
| BRAF | c.1799T>C | p.V600A | 7 | 140453136 | 140453136 |
| BRAF | c.1799T>G | p.V600G | 7 | 140453136 | 140453136 |
| BRAF | c.1798G>A | p.V600M | 7 | 140453137 | 140453137 |
| BRAF | c.1790T>G | p.L597R | 7 | 140453145 | 140453145 |
| BRAF | c.1790T>A | p.L597Q | 7 | 140453145 | 140453145 |
| BRAF | c.1789_1790CT>TC | p.L597S | 7 | 140453145 | 140453146 |
| BRAF | c.1789C>G | p.L597V | 7 | 140453146 | 140453146 |
| BRAF | c.1786G>C | p.G596R | 7 | 140453149 | 140453149 |
| BRAF | c.1781A>G | p.D594G | 7 | 140453154 | 140453154 |
| BRAF | c.1780G>A | p.D594N | 7 | 140453155 | 140453155 |
| BRAF | c.1406G>C | p.G469A | 7 | 140481402 | 140481402 |
| BRAF | c.1406G>T | p.G469V | 7 | 140481402 | 140481402 |
| BRAF | c.1406G>A | p.G469E | 7 | 140481402 | 140481402 |
| BRAF | c.1405G>A | p.G469R | 7 | 140481403 | 140481403 |
| BRAF | c.1397G>T | p.G466V | 7 | 140481411 | 140481411 |
| BRAF | c.1397G>A | p.G466E | 7 | 140481411 | 140481411 |
| EGFR | c.865G>A | p.A289T | 7 | 55221821 | 55221821 |
| EGFR | c.866C>T | p.A289V | 7 | 55221822 | 55221822 |
| EGFR | c.1793G>T | p.G598V | 7 | 55233043 | 55233043 |
| EGFR | c.2125G>A | p.E709K | 7 | 55241677 | 55241677 |
| EGFR | c.2126A>C | p.E709A | 7 | 55241678 | 55241678 |
| EGFR | c.2155G>A | p.G719S | 7 | 55241707 | 55241707 |
| EGFR | c.2155G>T | p.G719C | 7 | 55241707 | 55241707 |
| EGFR | c.2156G>C | p.G719A | 7 | 55241708 | 55241708 |
| EGFR | c.2235_2249del15 | p.E746_A750delELREA | 7 | 55242465 | 55242479 |
| EGFR | c.2236_2250del15 | p.E746_A750delELREA | 7 | 55242466 | 55242480 |
| EGFR | c.2237_2255>T | p.E746_S752>V | 7 | 55242467 | 55242485 |
| EGFR | c.2237_2251del15 | p.E746_T751>A | 7 | 55242467 | 55242481 |
| EGFR | c.2239_2248TTAAGAGAAG>C | p.L747_A750>P | 7 | 55242469 | 55242478 |
| EGFR | c.2239_2256del18 | p.L747_S752delLREATS | 7 | 55242469 | 55242486 |
| EGFR | c.2239_2251>C | p.L747_T751>P | 7 | 55242469 | 55242481 |
| EGFR | c.2239_2253del15 | p.L747_T751delLREAT | 7 | 55242469 | 55242483 |
| EGFR | c.2239_2247delTTAAGAGAA | p.L747_E749delLRE | 7 | 55242469 | 55242477 |
| EGFR | c.2240_2257del18 | p.L747_P753>S | 7 | 55242470 | 55242487 |
| EGFR | c.2240_2254del15 | p.L747_T751delLREAT | 7 | 55242470 | 55242484 |
| EGFR | c.2240T>C | p.L747S | 7 | 55242470 | 55242470 |
| EGFR | c.2254_2277del24 | p.S752_I759delSPKANKEI | 7 | 55242484 | 55242507 |
| EGFR | c.2290_2291ins12 | p.A763_Y764insFQEA | 7 | 55248992 | 55248993 |
| EGFR | c.2303G>T | p.S768I | 7 | 55249005 | 55249005 |
| EGFR | c.2307_2308insGCCAGCGTG | p.V769_D770insASV | 7 | 55249009 | 55249010 |
| EGFR | c.2308_2309insCCAGCGTGG | p.V769_D770insASV | 7 | 55249010 | 55249011 |
| EGFR | c.2311_2312insGCGTGGACA | p.D770_N771insSVD | 7 | 55249013 | 55249014 |
| EGFR | c.2319_2320insAACCCCCAC | p.H773_V774insNPH | 7 | 55249021 | 55249022 |
| EGFR | c.2369C>T | p.T790M | 7 | 55249071 | 55249071 |
| EGFR | c.2497T>G | p.L833V | 7 | 55259439 | 55259439 |
| EGFR | c.2504A>T | p.H835L | 7 | 55259446 | 55259446 |
| EGFR | c.2543C>T | p.P848L | 7 | 55259485 | 55259485 |
| EGFR | c.2573T>G | p.L858R | 7 | 55259515 | 55259515 |
| EGFR | c.2582T>A | p.L861Q | 7 | 55259524 | 55259524 |
| ERBB2 | c.929C>T | p.S310F | 17 | 37868208 | 37868208 |
| ERBB2 | c.1963A>G | p.I655V | 17 | 37879588 | 37879588 |
| ERBB2 | c.2033G>A | p.R678Q | 17 | 37879658 | 37879658 |
| ERBB2 | c.2264T>C | p.L755S | 17 | 37880220 | 37880220 |
| ERBB2 | c.2324_2325ins12 | p.A775_G776insYVMA | 17 | 37880995 | 37880996 |
| ERBB2 | c.2325_2326ins12 | p.A775_G776insYVMA | 17 | 37880996 | 37880997 |
| ERBB2 | c.2524G>A | p.V842I | 17 | 37881332 | 37881332 |
| KIT | c.1255_1257delGAC | p.D419del | 4 | 55589773 | 55589775 |
| KIT | c.1509_1510insGCCTAT | p.Y503_F504insAY | 4 | 55592185 | 55592186 |
| KIT | c.1669_1674delTGGAAG | p.W557_K558del | 4 | 55593603 | 55593608 |
| KIT | c.1669T>G | p.W557G | 4 | 55593603 | 55593603 |
| KIT | c.1669T>C | p.W557R | 4 | 55593603 | 55593603 |
| KIT | c.1669T>A | p.W557R | 4 | 55593603 | 55593603 |
| KIT | c.1669_1683del15 | p.W557_E561del | 4 | 55593603 | 55593617 |
| KIT | c.1670_1675delGGAAGG | p.W557_V559>F | 4 | 55593604 | 55593609 |
| KIT | c.1675_1677delGTT | p.V559del | 4 | 55593609 | 55593611 |
| KIT | c.1676T>A | p.V559D | 4 | 55593610 | 55593610 |
| KIT | c.1676T>C | p.V559A | 4 | 55593610 | 55593610 |
| KIT | c.1676T>G | p.V559G | 4 | 55593610 | 55593610 |
| KIT | c.1678_1680delGTT | p.V560del | 4 | 55593612 | 55593614 |
| KIT | c.1679T>A | p.V560D | 4 | 55593613 | 55593613 |
| KIT | c.1679T>G | p.V560G | 4 | 55593613 | 55593613 |
| KIT | c.1727T>C | p.L576P | 4 | 55593661 | 55593661 |
| KIT | c.1735_1737delGAT | p.D579del | 4 | 55593669 | 55593671 |
| KIT | c.1924A>G | p.K642E | 4 | 55594221 | 55594221 |
| KIT | c.1961T>C | p.V654A | 4 | 55594258 | 55594258 |
| KIT | c.2009C>T | p.T670I | 4 | 55595519 | 55595519 |
| KIT | c.2387G>A | p.R796K | 4 | 55599261 | 55599261 |
| KIT | c.2446G>T | p.D816Y | 4 | 55599320 | 55599320 |
| KIT | c.2446G>C | p.D816H | 4 | 55599320 | 55599320 |
| KIT | c.2447A>T | p.D816V | 4 | 55599321 | 55599321 |
| KIT | c.2466T>G | p.N822K | 4 | 55599340 | 55599340 |
| KIT | c.2466T>A | p.N822K | 4 | 55599340 | 55599340 |
| KIT | c.2474T>C | p.V825A | 4 | 55599348 | 55599348 |
| KIT | c.2485G>C | p.A829P | 4 | 55602664 | 55602664 |
| KRAS | c.437C>T | p.A146V | 12 | 25378561 | 25378561 |
| KRAS | c.436G>A | p.A146T | 12 | 25378562 | 25378562 |
| KRAS | c.351A>T | p.K117N | 12 | 25378647 | 25378647 |
| KRAS | c.351A>C | p.K117N | 12 | 25378647 | 25378647 |
| KRAS | c.183A>C | p.Q61H | 12 | 25380275 | 25380275 |
| KRAS | c.183A>T | p.Q61H | 12 | 25380275 | 25380275 |
| KRAS | c.182A>T | p.Q61L | 12 | 25380276 | 25380276 |
| KRAS | c.182A>G | p.Q61R | 12 | 25380276 | 25380276 |
| KRAS | c.182A>C | p.Q61P | 12 | 25380276 | 25380276 |
| KRAS | c.181C>A | p.Q61K | 12 | 25380277 | 25380277 |
| KRAS | c.181C>G | p.Q61E | 12 | 25380277 | 25380277 |
| KRAS | c.175G>A | p.A59T | 12 | 25380283 | 25380283 |
| KRAS | c.64C>A | p.Q22K | 12 | 25398255 | 25398255 |
| KRAS | c.57G>T | p.L19F | 12 | 25398262 | 25398262 |
| KRAS | c.40G>A | p.V14I | 12 | 25398279 | 25398279 |
| KRAS | c.38_39GC>AT | p.G13D | 12 | 25398280 | 25398281 |
| KRAS | c.38G>A | p.G13D | 12 | 25398281 | 25398281 |
| KRAS | c.38G>C | p.G13A | 12 | 25398281 | 25398281 |
| KRAS | c.38G>T | p.G13V | 12 | 25398281 | 25398281 |
| KRAS | c.37G>T | p.G13C | 12 | 25398282 | 25398282 |
| KRAS | c.37G>A | p.G13S | 12 | 25398282 | 25398282 |
| KRAS | c.37G>C | p.G13R | 12 | 25398282 | 25398282 |
| KRAS | c.35G>A | p.G12D | 12 | 25398284 | 25398284 |
| KRAS | c.35G>T | p.G12V | 12 | 25398284 | 25398284 |
| KRAS | c.35G>C | p.G12A | 12 | 25398284 | 25398284 |
| KRAS | c.34_35GG>TT | p.G12F | 12 | 25398284 | 25398285 |
| KRAS | c.34_35GG>CT | p.G12L | 12 | 25398284 | 25398285 |
| KRAS | c.34G>T | p.G12C | 12 | 25398285 | 25398285 |
| KRAS | c.34G>A | p.G12S | 12 | 25398285 | 25398285 |
| KRAS | c.34G>C | p.G12R | 12 | 25398285 | 25398285 |
| MET | c.1124A>G | p.N375S | 7 | 116340262 | 116340262 |
| MET | c.3029C>T | p.T1010I | 7 | 116411990 | 116411990 |
| MET | MET exon 14 skipping | MET exon 14 skipping | 7 | 116412042 | 116412042 |
| MET | MET exon 14 skipping | MET exon 14 skipping | 7 | 116412043 | 116412043 |
| MET | MET exon 14 skipping | MET exon 14 skipping | 7 | 116412044 | 116412044 |
| MET | MET exon 14 skipping | MET exon 14 skipping | 7 | 116412045 | 116412045 |
| MET | c.3757T>G | p.Y1253D | 7 | 116423428 | 116423428 |
| MET | c.3803T>C | p.M1268T | 7 | 116423474 | 116423474 |
| NRAS | c.183A>T | p.Q61H | 1 | 115256528 | 115256528 |
| NRAS | c.183A>C | p.Q61H | 1 | 115256528 | 115256528 |
| NRAS | c.182A>G | p.Q61R | 1 | 115256529 | 115256529 |
| NRAS | c.182A>T | p.Q61L | 1 | 115256529 | 115256529 |
| NRAS | c.182A>C | p.Q61P | 1 | 115256529 | 115256529 |
| NRAS | c.181C>A | p.Q61K | 1 | 115256530 | 115256530 |
| NRAS | c.181C>G | p.Q61E | 1 | 115256530 | 115256530 |
| NRAS | c.52G>A | p.A18T | 1 | 115258730 | 115258730 |
| NRAS | c.38G>A | p.G13D | 1 | 115258744 | 115258744 |
| NRAS | c.38G>T | p.G13V | 1 | 115258744 | 115258744 |
| NRAS | c.38G>C | p.G13A | 1 | 115258744 | 115258744 |
| NRAS | c.37G>C | p.G13R | 1 | 115258745 | 115258745 |
| NRAS | c.37G>T | p.G13C | 1 | 115258745 | 115258745 |
| NRAS | c.35G>A | p.G12D | 1 | 115258747 | 115258747 |
| NRAS | c.35G>T | p.G12V | 1 | 115258747 | 115258747 |
| NRAS | c.35G>C | p.G12A | 1 | 115258747 | 115258747 |
| NRAS | c.34G>A | p.G12S | 1 | 115258748 | 115258748 |
| NRAS | c.34G>T | p.G12C | 1 | 115258748 | 115258748 |
| NRAS | c.34G>C | p.G12R | 1 | 115258748 | 115258748 |
| PIK3CA | c.G241A | p.E81K | 3 | 178916854 | 178916854 |
| PIK3CA | c.263G>A | p.R88Q | 3 | 178916876 | 178916876 |
| PIK3CA | c.C277T | p.R93W | 3 | 178916890 | 178916890 |
| PIK3CA | c.G317T | p.G106V | 3 | 178916930 | 178916930 |
| PIK3CA | c.G323A | p.R108H | 3 | 178916936 | 178916936 |
| PIK3CA | c.A331G | p.K111E | 3 | 178916944 | 178916944 |
| PIK3CA | c.G353A | p.G118D | 3 | 178917478 | 178917478 |
| PIK3CA | c.1035T>A | p.N345K | 3 | 178921553 | 178921553 |
| PIK3CA | c.1258T>C | p.C420R | 3 | 178927980 | 178927980 |
| PIK3CA | c.1616C>G | p.P539R | 3 | 178936074 | 178936074 |
| PIK3CA | c.1624G>A | p.E542K | 3 | 178936082 | 178936082 |
| PIK3CA | c.1633G>A | p.E545K | 3 | 178936091 | 178936091 |
| PIK3CA | c.1633G>C | p.E545Q | 3 | 178936091 | 178936091 |
| PIK3CA | c.1634A>C | p.E545A | 3 | 178936092 | 178936092 |
| PIK3CA | c.1634A>G | p.E545G | 3 | 178936092 | 178936092 |
| PIK3CA | c.1635G>T | p.E545D | 3 | 178936093 | 178936093 |
| PIK3CA | c.1635G>C | p.E545D | 3 | 178936093 | 178936093 |
| PIK3CA | c.1636C>A | p.Q546K | 3 | 178936094 | 178936094 |
| PIK3CA | c.1636C>G | p.Q546E | 3 | 178936094 | 178936094 |
| PIK3CA | c.1637A>G | p.Q546R | 3 | 178936095 | 178936095 |
| PIK3CA | c.1637A>C | p.Q546P | 3 | 178936095 | 178936095 |
| PIK3CA | c.G1639A | p.E547K | 3 | 178936097 | 178936097 |
| PIK3CA | c.3062A>G | p.Y1021C | 3 | 178952007 | 178952007 |
| PIK3CA | c.3073A>G | p.T1025A | 3 | 178952018 | 178952018 |
| PIK3CA | c.3127A>G | p.M1043V | 3 | 178952072 | 178952072 |
| PIK3CA | c.3129G>T | p.M1043I | 3 | 178952074 | 178952074 |
| PIK3CA | c.G3129A | p.M1043I | 3 | 178952074 | 178952074 |
| PIK3CA | c.T3132A | p.N1044K | 3 | 178952077 | 178952077 |
| PIK3CA | c.3139C>T | p.H1047Y | 3 | 178952084 | 178952084 |
| PIK3CA | c.3140A>G | p.H1047R | 3 | 178952085 | 178952085 |
| PIK3CA | c.3140A>T | p.H1047L | 3 | 178952085 | 178952085 |
| PIK3CA | c.3145G>C | p.G1049R | 3 | 178952090 | 178952090 |
| PIK3CA | c.3145G>A | p.G1049S | 3 | 178952090 | 178952090 |

## Acknowledgments

We are very grateful to GenomiCare Biotechnology Co. Ltd., including data analysis by members of the bioinformatics department, testing work by the lab members, clinical data collection and entry by members of the clinical data department, and the collection and collation of patient follow-up information by members of the customer relationship department.

## Author contributions

Qiang Xu, Yanhong Deng, and Jiaping Li conceptualized and designed the study; Wei Jiang and Jinwang Wei contributed to the experimental design and methodology, and administered the project; Jianwei Zhang, Linbo Cai, Minjie Luo, and Zhan Wang communicated with patients, and collected and organized patient follow-up information; Chen Wang, Wending Sun, and Chun Dai performed the experimental work; Jinwang Wei, Guan Wang, and Qiang Xu performed the bioinformatics and statistical analysis; Wending Sun wrote the manuscript. All authors approved the final version of the manuscript.

## Data and materials availability

The datasets used and/or analyzed in the study are available from the corresponding author on reasonable request.

